# Biomolecular Tau condensation is linked to Tau accumulation at the nuclear envelope

**DOI:** 10.1101/2022.01.24.477544

**Authors:** Janine Hochmair, Christian Exner, Maximilian Franck, Alvaro Dominguez-Baquero, Lisa Diez, Hévila Brognaro, Matthew Kraushar, Thorsten Mielke, Helena Radbruch, Senthil Kaniyappan, Sven Falke, Eckhard Mandelkow, Christian Betzel, Susanne Wegmann

## Abstract

Biomolecular condensation of the neuronal microtubule-associated protein Tau (MAPT) can be induced by coacervation with polyanions like RNA, or by molecular crowding. Tau condensates have been linked to both functional microtubule binding and pathological aggregation in neurodegenerative diseases. We find that molecular crowding and coacervation with RNA, likely coexisting in the cytosol, synergize to enable Tau condensation at physiological buffer conditions and produce condensates with a strong affinity to charged surfaces. During condensate-mediated microtubule polymerization, this synergy enhances bundling and spatially arranges microtubules. We further show that different Tau condensates efficiently induce pathological Tau in cells, including small accumulations at the nuclear envelope that correlate with nucleocytoplasmic transport deficits. Fluorescent lifetime imaging reveals different molecular packing densities of Tau in cellular accumulations, and a condensate-like density for nuclear envelope Tau. These findings suggest that a complex interplay between interaction partners, post-translational modifications, and molecular crowding regulates the formation and function of Tau condensates. Conditions leading to prolonged existence of Tau condensates may induce the formation of seeding-competent Tau and lead to distinct cellular Tau accumulations.

## Introduction

The intrinsically disorded protein Tau is a neuronal microtubule (MT) binding protein that helps to stabilize axons in the central nervous system. In Alzheimer’s disease (AD) and other tauopathies, Tau accumulates intraneuronally and aggregates as neurofibrillary tangles (NFTs) into highly ordered β-structured Tau aggregates, which are a hallmark of the disease and are associated with progressive neuronal death (Braak & Braak, 1991). Tau aggregation and spreading in the brain are thought to involve the seeding of Tau aggregation by small amounts of misfolded Tau (Dujardin & Hyman, 2019; Frost & Diamond, 2010), a mechanism described as ‘templated misfolding’ or ‘seeded aggregation’. Interestingly, the morphology and molecular content of intraneuronal Tau aggregates in the brain is variable and disease specific (Ferrer *et al*, 2014), and also more subtle forms of Tau accumulations, for example around the nucleus, can occur (Eftekharzadeh *et al*, 2018; Cornelison *et al*, 2019). Tau accumulating at the nuclear envelope was shown to inhibit neuronal nucleocytoplasmic transport of proteins (Eftekharzadeh *et al*, 2018) and RNA (Cornelison *et al*, 2019), induce changes in nuclear morphology (Sheffield *et al*, 2006) and lamin folds (Frost *et al*, 2016), and leads to MTs invading the nucleoplasm (Paonessa *et al*, 2019). The molecular assembly state of Tau at the nuclear envelope is not known, but it appears that they differ from amyloid-like large cytosolic aggregates since they have never been reported by investigations of Tau pathology using amyloid-dyes. Cell models of seeded Tau aggregation can in part recapitulate different types of Tau accumulations (Holmes *et al*, 2014), whereby the morphology and subcelluar localization of the accumulations is goverened by the Tau material used as seeds (Kaufman *et al*, 2016), and the molecular packing of Tau in various described accumulations seems to differ (Kaniyappan *et al*, 2020).

Tau can have different assembly forms that all display unique biophysical and biochemical characteristics, and have different cellular (mis)functions. Monomers and dimers are widely thought to resemble the soluble ‘native’ form of Tau in the cytosol. Tau oligomers seem to encode neurotoxicity (Ward *et al*, 2012) and are discussed to seed aggregation and mediate Tau pathology spreading between neurons (Gerson & Kayed, 2013). Tau aggregates rich in β-structure are the stable end product of Tau aggegation processes and are deposited in long-lasting neuronal inclusions in the brain (Brion, 1998). In addition, liquid-like protein condensates of Tau have been suggested to be involved in different Tau functions, including the binding and polymerization of MTs (Hernández-Vega *et al*, 2017; Siahaan *et al*, 2019; Tan *et al*, 2019; Zhang *et al*, 2020) as well as the *de novo* formation of seeding competent Tau oligomers (Wegmann *et al*, 2018; Boyko *et al*, 2020; Kanaan *et al*, 2020).

Liquid-like condensates of Tau form via liquid-liquid phase separation (LLPS), a process that is used by cells to assemble membrane-less organelles of various functions (Mitrea & Kriwacki, 2016; Hyman *et al*, 2014). From studies *in vitro*, two different modes of Tau condensation have been identified. Tau LLPS can be initiated through the addition of macromolecular cowding agents like polyethylene glycol (PEG) or dextrane to a dilute Tau protein solution (Wegmann *et al*, 2018; Ambadipudi *et al*, 2017). The crowding-induced colloid osmotic pressure (Mitchison, 2019), together with attractive forces between Tau molecules, triggers the de-mixing of Tau into liquid-dense condensates. Crowding-induced biomolecular Tau condensates have been suggested to harbor pathological seeding potential since they can convert - at least *in vitro* - into oligomeric Tau species with properties similar to *in vitro* generated Tau aggregates (Boyko *et al*, 2020; Kanaan *et al*, 2020; Wegmann *et al*, 2018).

Tau LLPS can also be induced through complex coacervation, which is the spontaneous organization of oppositely charged (parts of) biomolecules into higher order assemblies, for example liquid-like condensates. This process is driven by electrostatic interactions of the engaging molecules. In the case of Tau, polyanionic RNA (or heparin (Lin *et al*, 2020)) co-condensate with the positively charged Tau microtubule-assembly domain into liquid-like droplets (Zhang *et al*, 2017; Lin *et al*, 2019). It is not known whether Tau:RNA coacervates can trigger the formation of pathological Tau aggregates similar to crowding-induced Tau condensates. Notably, high concentrations of RNA do not induce Tau LLPS but can trigger Tau aggregation (Kampers *et al*, 1996) without the formation of liquid-condensed intermediates (Lin *et al*, 2020). In cells, Tau coacervation may, for example, be important for the co-condensation of Tau with RNA-containing ribonucleoparticles such as stress granules (Vanderweyde *et al*, 2016; Ash *et al*, 2021), as well as for the binding of Tau to the anionic MT surface (Mukrasch *et al*, 2007; Kadavath *et al*, 2018). It has been suggested that the binding of Tau to MTs involves both single Tau molecules (Janning *et al*, 2014) as well as liquid-condensed forms of Tau (Hernández-Vega *et al*, 2017; Tan *et al*, 2019; reviewed in (Mitchison, 2020)). In the following, we will use ‘*condensation’* to describe all LLPS processes including those induced by crowding, polyanions, or both. The term ‘*coacervation’* refers solely to the condensation of Tau with polyanionic co-factors, in the absence of crowding.

In the neuronal cytoplasm, Tau encounters RNA, microtubules, and other polyanions, and thus may undergo crowding- and polyanion-induced condensation at the same time. To mimic this scenario, we characterized Tau LLPS in model systems that consider the contribution of both crowding and polyanions. We find that, in fact, molecular crowding is necessaryto enable Tau and phosphorylated Tau coacervation with RNA at intracellular ion compositions. The Tau binding partners RNA and tubulin seem to compete for co-condensation with Tau, and the presence of RNA during MT polymerization out of Tau condensates enables pronounced MT bundling and a unique geometric bundle arrangement, likely a result of the high binding affinity (wetting) to negatively charged MT surfaces combined with strong cohesive forces inside of Tau:PEG:RNA condensates.The generation of seeding-competent Tau is similar for Tau condensates with PEG, with RNA, or with both, and all aged Tau condensates convert into similar types of cytosolic and nuclear Tau accumulations, including Tau at the nuclear envelope. Combining FRAP and FLIM of CFP-Tau, we find that different types of Tau accumulations display modes of molecular packing reminiscent of cytosolic and nuclear aggregates versus less densely packed condensates at the nuclear envelope. These accumulations further enhance Tau-induced nucleocytoplasmic transport deficits present in Tau expressing cells. Interestingly, binary Tau:RNA coacervates promote Tau seed formation in the absence of condensate liquid-to-gel transition, which deviates from the current model of progressive percolation formulated for RNA-binding protein condensates.

Collectively, these observations provide a framework to explain the formation, interactions, functions, and pathological activities of different biomolecular Tau condensates, and thereby improve the understandig of how condensation contributes to Tau biology and disease-related cellular Tau accumulations.

## Results

### RNA and crowding synergize during Tau condensation

Tau condensation *in vitro* can be induced by molecular crowding agents or by polyanions like RNA. The cell has a generally crowded environment, and RNA may be a potent inducer of Tau in the cytosol. Cellular Tau condennsation may thus be influenced by both factors at the sam time. To test whether Tau condensates could form in an environment where both RNA and molecular crowding are present, we mixed human full-length wildtype Tau (5 µM Tau, 2N4R isoform, 441 aa, net charge at pH 7.4 = +1.4 (unphosphorylated); Figure 1A) with polyA RNA at charge matching concentrations, and with the molecular crowding agent polyethylene glycol (PEG8000; 5% (w/vol)). Light microscopy confirmed the previously reported formation of liquid-like Tau phases (observed as droplets) in the presence of RNA (Tau:RNA) or PEG (Tau:PEG), as well as in tertiary systems containing both polyA and PEG (Tau:PEG:RNA; Figure 1B). Using fluorescently labeled Tau-Dylight488, polyA-Cy3, and PEG-Dylight647, we observed that Tau and RNA become enriched inside the condensates, whereas the crowding agent PEG is excluded (Figure 1C).

**Figure 1.**
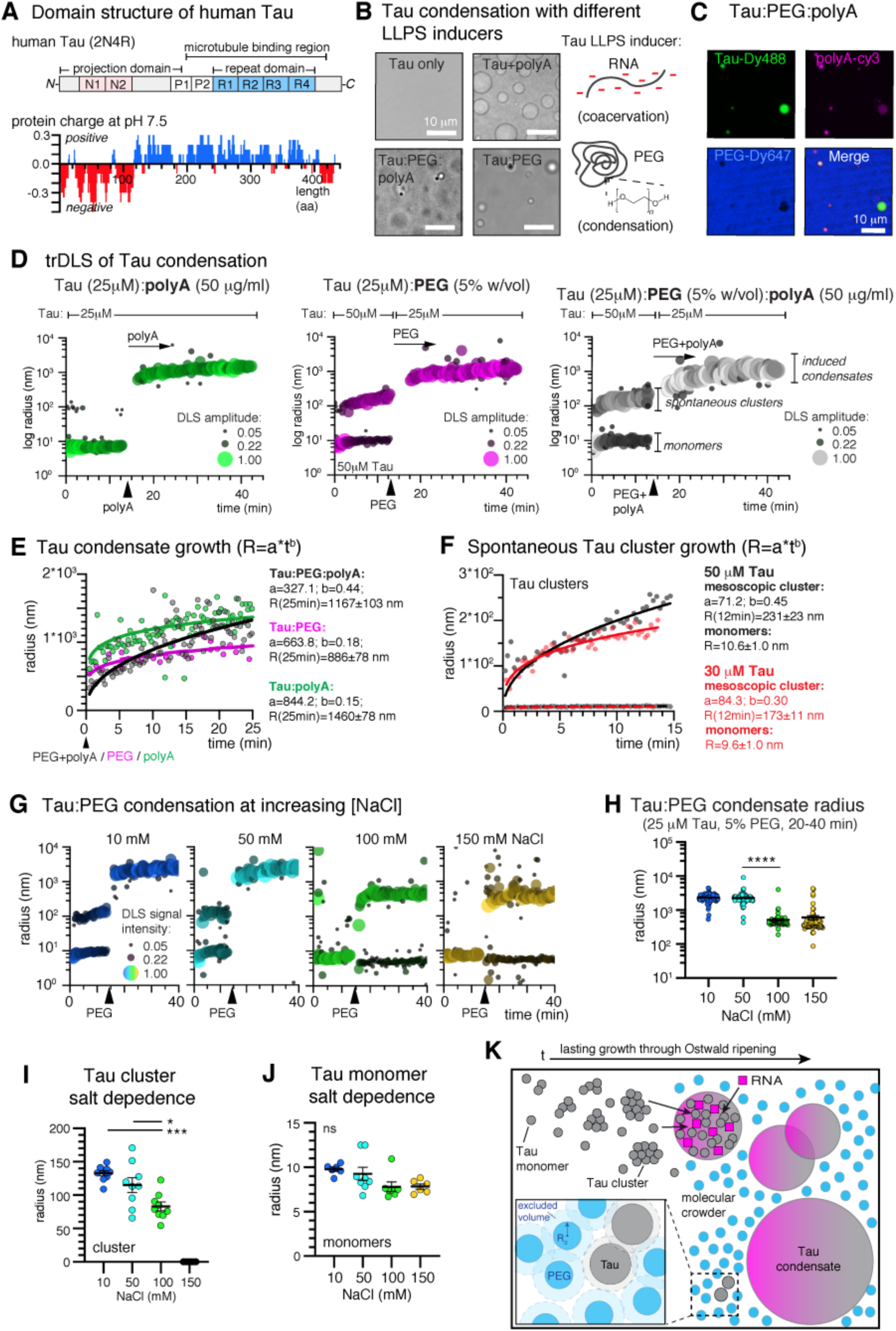
Kinetics of Tau condensation and mesoscopic cluster formation. A Domain structure of the longest human Tau isoform (2N4R) used in this study: the positively charged C-terminal half of Tau contains four repeats (R1-R4) and two proline rich regions (P1, P2), and comprises the microtubule binding region of Tau. This region binds to polyanionic macromolecules such as RNA through electrostatic interactions. The N-terminal ∼200 amino acids (aa) project from the microtubule surface and contain two negatively charged N-terminal inserts (N1, N2). The homogenous charge distribution (sliding window of 10 aa) along its aa sequence assigns Tau an amphiphilic character. B Addition of polyA RNA, PEG (PEG8000), or both induces the de-mixing of Tau at physiological concentrations (5 μM Tau in 25 mM HEPES, 1 mM DTT, pH 7.4). Scale bars=10 μm. C Confocal fluorescence microscopy shows co-condensation of polyA-Cy3 (10:1, polyA:polyA-Cy3) with Tau (5 µM Tau with 1% Tau-DyLight488). PEG8000-Dylight647 is excluded from Tau condensates. Scale bar=10 μm. D Size distribution of molecular Tau assemblies during Tau LLPS measured by time-resolved DLS (trDLS). At 25 μM Tau in low salt (10 mM NaCl) buffer, trDLS detects Tau monomers (radii ∼10 nm) and mesoscopic Tau clusters (radii ∼100-200 nm). Addition of PEG, polyA, or both after ∼15 min induced the formation of Tau condensates with a radius of >10^3^ nm, at the cost of monomers and mesoscopic clusters in the solution. Data point bubble sizes correspond to DLS amplitudes, which are proportional to the intensity of scattered light of the respective particles. E Kinetics of Tau:PEG, Tau:RNA, and Tau:PEG:RNA condensate growth assessed through power law, R=a*t^b^, fitting to trDLS data starting at the time of LLPS inducer addition. F Kinetics of cluster growth in low salt (25 mM HEPES, 10 mM NaCl) at 30 μM and 50 μM Tau using the same fitting model as in (E). G trDLS of Tau:PEG condensation at increasing NaCl concentrations. H Radius of Tau:PEG condensates at different NaCl concentrations. Data shown as mean±SEM. I Radius of Tau clusters at different NaCl concentrations. Data shown as mean±SEM. J Radius of Tau monomers at different NaCl concentrations. Data shown as mean±SEM. K Model of Tau condensation: Tau monomers (grey circles) and clusters coacervate with RNA (pink) to form condensates that grow through droplet fusion (Oswald ripening). Molecular crowding agents, like PEG (blue), are excluded from the dense phase. Inset: excluded volume effects exerted by PEG force individual Tau molecules to interact, thereby trigger LLPS and stabilize Tau:RNA coacervates against the electrostatic shielding of molecular interactions inside the dense phase. As a result, Tau:PEG:RNA condensates can form at cytosol-like ion concentrations. R_0_=radius of hydration. Data in H, I, J have been compared by one-way ANOVA with Tukey test for multiple comparison.

To describe the effects of RNA and crowding on the Tau condensation process, we performed time-resolved dynamic light scattering (trDLS), which allows to follow the evolution and size distribution of particles in a particle radius (R_0_, radius of hydration) range between 10^0^ to 10^5^ nm in solution over time (Van Lindt *et al*, 2021; Falke *et al*, 2019; Schubert *et al*, 2017). During the first ∼15 min of measuring the particle radii in a Tau solution (25 μM for Tau:polyA, 50 μM for Tau:PEG and Tau:PEG:polyA), we observed Tau monomers (R_Mono_∼10 nm) and spontaneously formed Tau clusters (R_Clust_∼200 nm; Figure 1D; Expanded View Figure EV1A,B). Addition of polyA (50 μg/ml) or PEG (5%) induced an immediate increase in particle radius, which could be attributed to the formation of Tau condensates, as judged by light microscopy. Simultaneous addition of polyA and PEG induced a similar increase of particle radii in the solution. Notably, the trDLS measurements were recorded in low salt buffer (25 mM HEPES, pH7.4, 10 mM NaCl) to enable Tau:RNA coacervation.

Liquid-like protein condensates grow through the recruitment of monomers from the dilute phase and through droplet coalescence (Ostwald ripening; (Lifshitz & Slyozov, 1961; Siggia, 1979; Lee *et al*, 2021)), which can be described by the power law *R=a*t^b^*, with *R* being the condensate radius, *t* the time after LLPS initiation, and *b* the coarsening exponent describing the increase in particle sizes (Berry *et al*, 2018); *b*∼1/3 describes the growth by Ostwald ripening of most biomolecular condensates. The growth kinetics of Tau:PEG:RNA condensates (b=0.44; R_25min_: 1167±103 nm) indicated a major contribution of Ostwald ripening during their formation. Binary Tau:RNA and Tau:PEG condensates showed fast initial growth, which then slowed down (polyA: b=0.15; R_25min_: 1460±78 nm; PEG: b=0.18; R_25min_: 886±78 nm), indicating less contribution of Ostwald ripening compared to the triple LLPS system (Figure 1E). Tau:PEG systems produced the smallest condensates. We also induced Tau coacervation with other polyanions (heparin, tRNA, and polyU RNA) that were previously used to induce Tau LLPS, and found that all conditions resulted in similar Tau condensation responses (Expanded View Figure EV1C).

In summary, we found that polyanions and crowding synergize during Tau condensation producing Tau coacervates that exhibit growth kinetics as previously reported for other biomolecular condensates (Van Lindt *et al*, 2021).

### Mesoscopic Tau clusters below c_sat_ for microscopic condensation

The spontaneous formation of microscopic liquid condensates, i.e. protein droplets large enough to be identified as such by light microscopy, occurs when the protein concentration exceeds the critical saturation concentration, c_sat_ (Shin & Brangwynne, 2017). For Tau, spontaneous microscopic condensation, in the absence of cofactors, was previously observed by light microscopy at concentrations of ∼50 to 100 μM Tau (Wegmann *et al*, 2018; Ambadipudi *et al*, 2017). In our trDLS measurements, before the addition of RNA and PEG, we observed the formation of small spontaneous Tau clusters with a radius of ∼150 to 250 nm already at 25-50 μM Tau (Figure 1D). Size and formation kinetics of these clusters depended on the Tau concentration and followed kinetics similar to Tau coacervates, reminiscent of Ostwald ripening (30 μM Tau: R_12min=_173.0±10.7 nm, b=0.30; 50 μM Tau: R_12min_: 230.8±22.7 nm, b=0.45; Figure 1F). To test if Tau cluster formation depended on electrostatic interactions between and within Tau molecules, we performed trDLS of Tau condensation at different NaCl concentrations (Figure 1G). The size of Tau:PEG condensates (Figure 1H) and spontaneous Tau clusters (Figure 1I) both decreased with increasing salt concentration. Only few Tau clusters formed at 100 mM NaCl and none at 150 mM NaCl, indicating that spontaneously formed Tau clusters below c_sat_ depended on electrostatic interactions between Tau molecules. Notably, monomeric Tau had a radius of 9.6±1.0 nm (mean±SD) at 30 μM Tau and 10 mM NaCl, consistent with previous reports (Mylonas *et al*, 2008), which was rather insensitive to increasing ion concentrations (Figure 1J).

Tau clusters coexisted with Tau monomers and disappeared upon addition of RNA or PEG to induce microscopic LLPS (Figure 1D,G). We suggest that the amphiphilic character of Tau (Figure 1A) enables the formation of small (=mesoscopic) Tau clusters at c<c_sat_, therefore preceding microscopic LLPS (Figure 1K).

In summary, we identified mesoscopic Tau clusters that form spontaneously in solution based on electrostatic interactions and coexist with monomeric Tau, but disappear in favor of larger microscopic liquid-like Tau condensates under conditions enabling LLPS *in vitro*. The mesoscopic Tau clusters could resemble condensate pre-cursors - not previously detected due to resolution limits of light microscopy - that present “emerging stickers” (Choi *et al*, 2020a) and thereby enable heterologous nucleation (Hyman *et al*, 2014) of Tau condensation, or nucleated crystal formation (Schönherr *et al*, 2015; Schubert *et al*, 2017). They may thus play an important role for functional and pathological LLPS of Tau. Recently, similar mesoscopic clusters were reported for p53 in cells (Yang *et al*, 2020).

### Crowding enables Tau coacervation at physiological ion concentrations

Tau coacervates assemble via electrostatic interactions between positive charges in Tau and negative charges in RNA molecules (Lin *et al*, 2019) or between oppositely charged Tau domains (Boyko *et al*, 2019), which infers a sensitivity to the ion concentration in the buffer system. Since the ion concentration of the cytosol exceeds that of low salt buffer systems commonly used in Tau coacervation studies, one may question the relevance of Tau coacervation in the cellular context. Therefore, we formed different Tau condensates at 10 mM NaCl and tested, to which degree they could withstand a step-wise increase in NaCl concentrations.

In the binary Tau:RNA system, condensates disappeared already at 25 mM NaCl (Figure 2A), confirming the high sensitivity to the shielding of electrostatic interactions in their interior by counter ions in the buffer. Tau:PEG condensates also formed at higher NaCl concentrations, whereby their size decreased at ≥100 mM NaCl (Figure 1G,H; Radius (mean±SEM) at 50 mM NaCl: 2264±108nm; at 100 mM: 515±47 nm; p<0.0001), accompanied by the re-occurrence of monomers. Tau:PEG:RNA condensates did not change between 5 and 150 mM NaCl, and smaller condensates still robustly formed at 300 mM NaCl levels (Figure 2A). In axoplasm-like ion composition (X/2 buffer (Song & Brady, 2013)), the formation of condensates occurred in Tau:PEG and more robustly (more and larger condensates) in Tau:PEG:RNA systems (Figure 2A,B,C). Tau:RNA coacervation was inhibited in X/2 buffer. These data suggested that crowding is necessary for Tau condensation at physiological salt concentrations.

**Figure 2.**
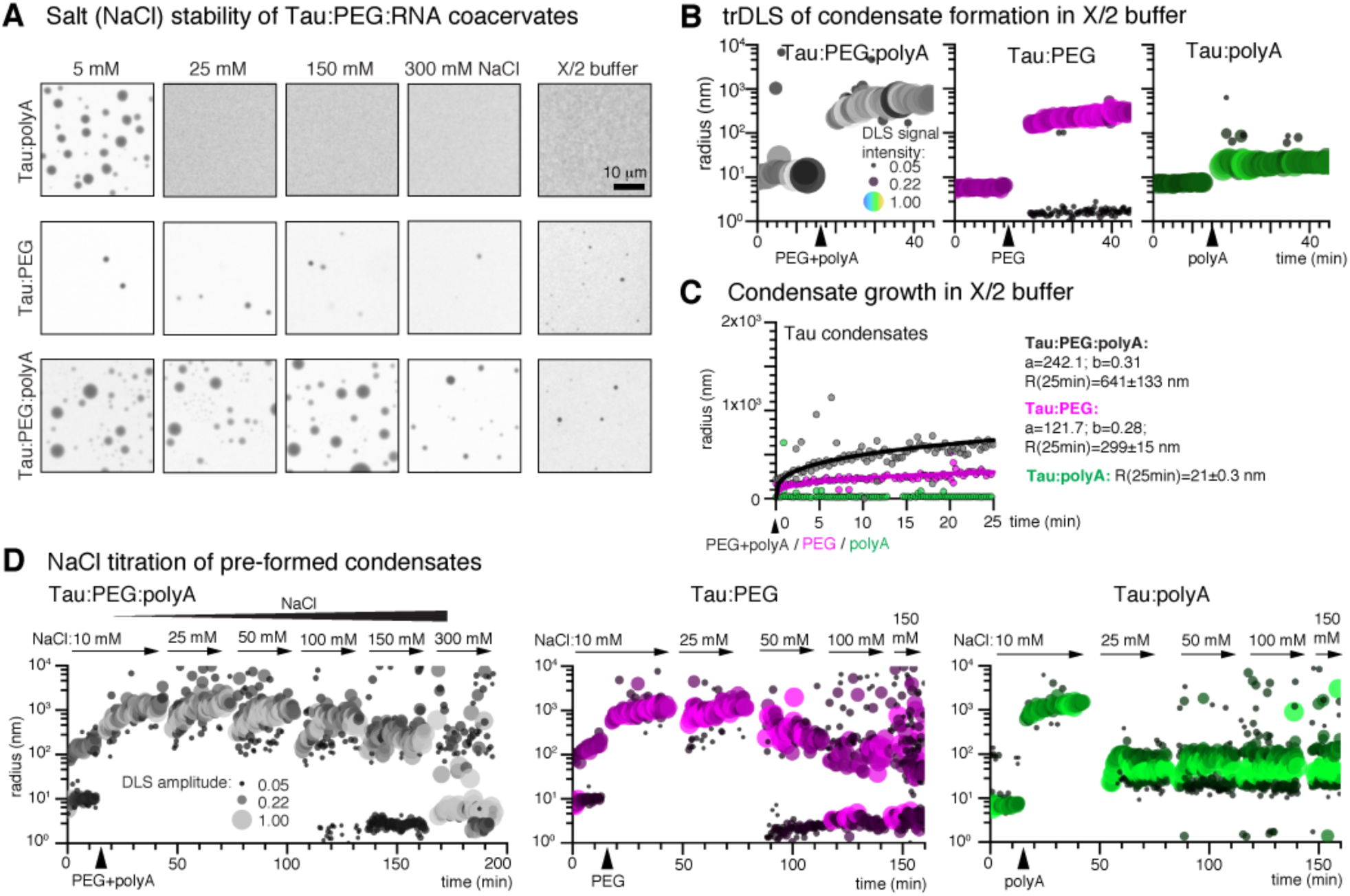
Ion strength dependence of Tau coacervation is overcome by crowding. A Microscopy of Tau:polyA, Tau:PEG, and Tau:PEG:polyA condensates (5 µM Tau, 10% PEG, 4.8 µg/ml polyA) formed at increasing NaCl concentrations and in X/2-buffer (Song & Brady, 2013) mimicking the axoplasm ion composition. Scale bar=10 μm. B trDLS of Tau condensation in X/2-buffer. C Condensate growth kinetics in X/2 buffer assessed through power law, R=a*t^b^, fitting to trDLS data starting at the time of LLPS inducer addition. D trDLS of Tau condensation in low salt buffer, with subsequent stepwise increase of NaCl. In B and C, data point bubble sizes correspond to DLS amplitudes.

We confirmed these results by trDLS, whereby we monitored Tau condensation at 10 mM NaCl and then step-wise increased the NaCl concentration. Whereas Tau:RNA coacervates disassembled already 25 mM NaCl, Tau:PEG condensates appeared destabilized at 100 mM NaCl, and Tau:PEG:RNA condensates at 300 mM NaCl (Figure 2D). Interestingly, similar experiments using phosphate instead of NaCl titration revealed a higher sensitivity of Tau:PEG and Tau:PEG:RNA condensates to phosphate ions. Tau:PEG:RNA condensates – pre-formed in 10 mM phosphate – disassembled in the presence of 100 mM phosphate, and Tau:PEG condensates already at 75 mM phosphate (Expanded View Figure EV2A,B). Furthermore, when Tau condensates disappeared, Tau clusters and monomers re-occurred, supporting our hypothesis that Tau clusters could be precursors of Tau condensates.

Together these data show that Tau:RNA coacervates, relying solely on electrostatic interactions, are easily inhibited by low amounts of counter ions and, thus, are not likely to form in cytosolic ion conditions. Crowding-induced condensates seem to include other interactions and are hence less sensitive to ion concentration. Furthermore, the interactions stabilizing Tau condensates appear to be more sensitive to phosphate compared to NaCl ions in the buffer, which may underlie a similar mechanism as the previously reported hydrotropic effect of adenosine triphosphate (ATP) on biomolecular condensates (Patel *et al*, 2017). For the condensation of Tau in cells this implicates that different Tau LLPS mechanisms need to synergize to form condensates that are stable at physiological ion composition, and that condensates may be modulated by changes in the (local) ion composition.

### Molecular crowding enables coacervation of phosphorylated Tau

Human Tau carries >80 putative phosphorylation sites in its amino acid sequence and is a substrate for multiple kinases and phosphatases, consequently leading to a complex condition-dependent mix of phospho-Tau variants in the cell. Phosphorylation of Tau changes its charge from positive (+1.5 at pH 7.4) to negative, which reverses Tau’s intrinsic cationic character towards neutral or anionic and, thus, may impact the capability of phospho-Tau to undergo polyanion-induced coacervation.

To test this idea, we performed LLPS experiments using p-Tau from Sf9 cells that presents an ensemble of phosphorylation states ((Drepper *et al*, 2020); ∼4-12 phosphates per Tau molecule; mean occupancy ∼8, net charge ∼10.5, assuming an average charge of -1.5 per phosphate). Indeed, microscopy (Figure 3A) and trDLS (Expanded View Figure EV2C) experiments supported our hypotheses: p-Tau failed to coacervate with RNA but the condensation could still be induced with PEG. In p-Tau:PEG:RNA systems, we could observe the co-partitioning of RNA into Tau condensates (Figure 3A), and both p-Tau:PEG:RNA and p-Tau:PEG condensates formed at elevated NaCl concentrations (Expanded View Figure EV2D). Since most, if not all, Tau in cells is phosphorylated under physiological conditions, molecular crowding seems essential to enable Tau condensation in the cellular environment.

**Figure 3.**
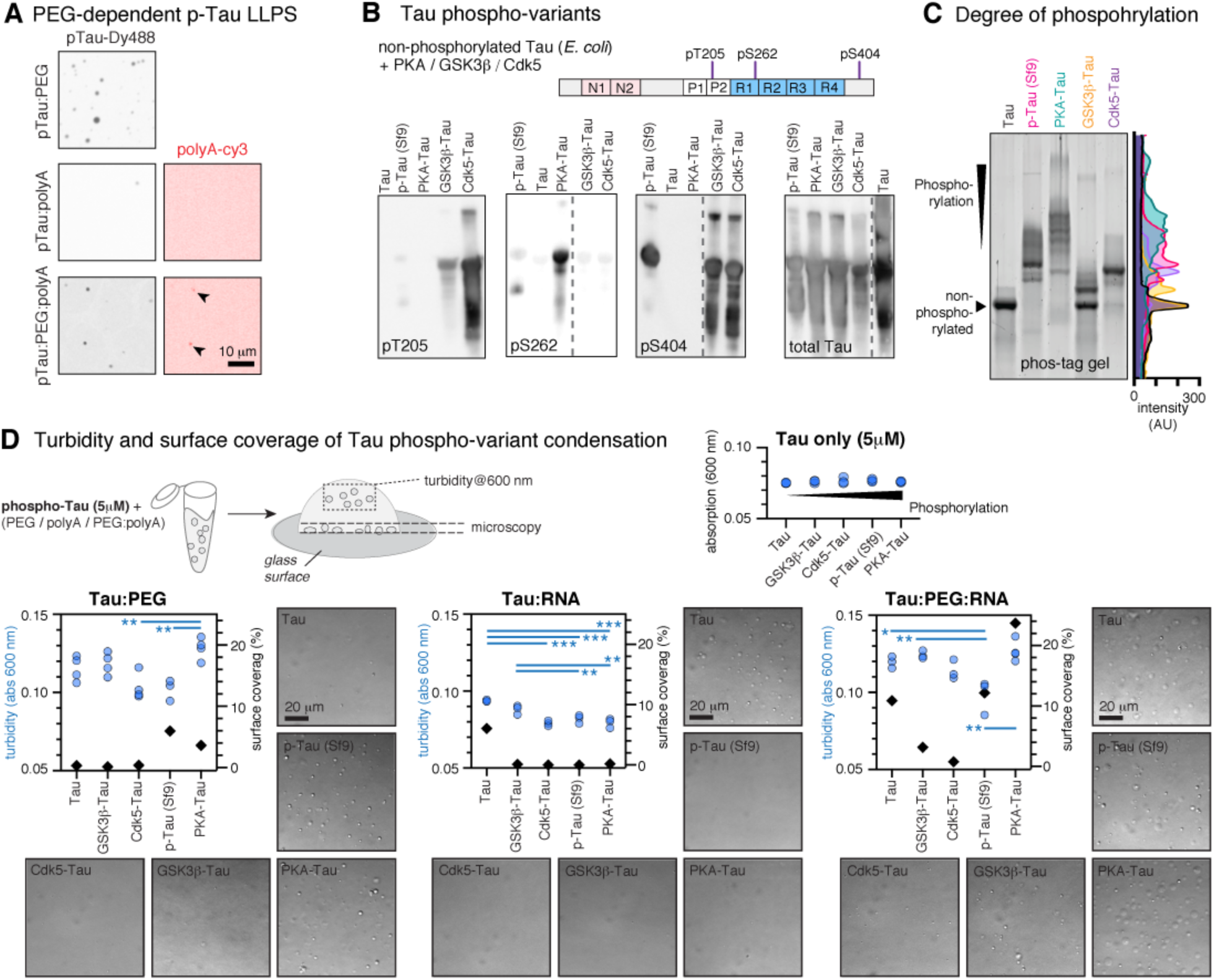
Phosphorylation modulates Tau condensation and condensate affinity to surfaces. A Recombinant phosphorylated Tau expressed in insect cells (p-Tau) de-mixes in the presence of PEG but fails to coacervate with RNA (5 μM p-Tau, 10% PEG). Coacervation with PEG and polyA-Cy3 shows co-condensation of RNA with p-Tau (arrow heads). Scale bar=10 μm. B Tau phospho-variants generated by *in vitro* phosphorylation of non-phosphorylated Tau from *E. coli* with different kinases (PKA, GSK3β, or Cdk5) show differences in phosphorylation at specific phospho-site (pT205, pS262, and pS404) tested by Western blot. Non-phosphorylated Tau and p-Tau from insect cells are loaded for comparison. Position of tested p-sites is indicated in the Tau sequence cartoon. Dotted vertical lines in Western blot images indicate that lanes are from different immunoblots. C Phos-tag gel shows degree and heterogeneity of phosphorylation in the different Tau phospho-variants. Intensity along the lanes are plotted as colored lines next to gel image. D Condensation of Tau phospho-variants in the Tau:PEG, Tau:RNA (polyA), and pTau:PEG:RNA systems was determined by turbidity measurements for condensates in solution above the glass surface (absorption at 600 nm; blue circles) and by microscopy for surface attached condensates (surface coverage in %; black symbols). One-way ANOVA with Tukey post-test. Scale bars=20 μm.

To further explore the effect of physiological Tau phosphorylation on condensation, we investigated Tau phospho-variants generated via *in vitro* phosphorylation of non-phosphorylated full-length Tau (2N4R; from E. coli) using kinases (GSK3β, PKA, and Cdk5) known to differentially phosphorylate Tau (Liu *et al*, 2006). The three kinases phosphorylated Tau at multiple and overlapping sites, yet produce differential phosphorylation at the Tau phosphorylation sites pT205, pS262, and pS404 (Figure 3B). Whereas pS404 is equally abundant in Cdk5-Tau and GSK3β- Tau, pT205 is abundant in Cdk5-Tau and less in GSK3β-Tau, and pS262 seems specific for PKA-Tau. p-Tau from insect cells is highly phosphorylated at pS404, and far less at pS262 and pT205. To estimate the overall amount of phosphorylation, we separated the Tau phospho-variants on phos-tag gels, in which the electrophoretic separation of proteins occurs according to the number of attached phospho-groups (Kinoshita *et al*, 2006; Samimi *et al*, 2021). All phospho-variants showed multiple bands, indicating the expected heterogenous degree of phosphorylation per molecule in each sample (Figure 3C). On average, Tau phosphorylated by PKA had the highest amount of phosphate groups, followed by p-Tau (from Sf9 cells) and Cdk5-phosphorylated Tau having a similar degree of phosphorylation. In case of Tau phosphorylated by GSK3β, about 50% of the molecules remained non-phosphorylated, and a second prominent phospho-species occurred, similar to previous observations (Liu *et al*, 2006).

We then tested the ability of these phospho-variants for undergoing LLPS in the presence of either PEG or RNA, or both. In our experiments, we observed that Tau condensates with a high affinity for charged surfaces successively deplete from the solution and become invisible to turbidity measurements. To assess the overall amount of condensation, we therefore determined the amount of condensates in solution through turbidity measurements (absorption at λ=600 nm) and in the same samples assessed the amount of condensates wetting the supporting glass surface (surface coverage in %) by microscopy (Figure 3D). LLPS of all phospho-variants was diminished in pTau:RNA systems, at least at the given RNA concentration used (0.05 μg/ml polyA). Non-phosphorylated Tau showed stronger surface affinity in Tau:RNA compared to Tau:PEG systems. In pTau:PEG systems, the amount of total LLPS (turbidity+surface coverage) was higher for p-Tau and PKA-Tau compared to non-phosphorylated Tau, and their high phosphorylation status increased the attachment of these condensates to the glass surface. LLPS of GSK3β-Tau and Cdk5-Tau, both highly phosphorylated at pS404 and pT205, was similar to non-phosphorylated Tau. In Tau:PEG:RNA systems, we observed the highest in-solution LLPS and surface wetting for PKA-Tau, whereas p-Tau behaved similar to non-phosphorylated Tau. This indicates that phosphorylation above a certain threshold may drive LLPS in this system, or that the combination of phospho-sites present in PKA-phosphorylated Tau but distinct from p-Tau (Drepper *et al*, 2020) may drive Tau LLPS. Interestingly, we previously found that Tau phosphorylated by MARK2 (main target sites pS262, pS356) also enhances PEG-induced Tau LLPS (Wegmann *et al*, 2018). In summary, we find that phosphorylation generally inhibits Tau coacervation with RNA, but in the presence of crowding and/or RNA, pS262 seems to favor Tau LLPS whereas pT205 and pS404 (and other p-sites) may have no effect or rather inhibit Tau condensation. Similarly, it was shown that pseudo-phosphorylation in the proline-rich domain of Tau, including at T205, can inhibit cellular Tau condensation (Zhang *et al*, 2020). Residues S262 and S404 are located outside of this region. These data show that physiological Tau phosphorylation can have an influence on Tau LLPS, and that certain phospho-sites may play a role in regulating cellular Tau condensation. However, due to the high complexity of Tau post-translational modifications, further investigations are needed to decipher the effects of individual, or groups of, PTMs on Tau LLPS.

### Tau coacervation modulates Tau:tubulin interactions

It was shown *in vitro* that crowding-induced Tau condensates can nucleate MTs through co-partitioning of tubulin (Hernández-Vega *et al*, 2017), and that Tau can form condensed ‘islands’ on individual MTs (Tan *et al*, 2019; Siahaan *et al*, 2019). In cells, Tau condensation appears to catalyze the binding to MTs (Zhang *et al*, 2020).

The initial step during condensate-mediated MT nucleation is the co-condensation of tubulin with Tau. To understand the influence of crowding and RNA on this process, we prepared Tau:PEG, Tau:RNA, and Tau:PEG:RNA condensates (25 μM Tau-Dylight488, 25 μg/ml polyA-Cy3, 5% PEG) and mixed them with tubulin (5 μM, 1:10 tubulin-HiLyte647). In all three LLPS systems, tubulin readily co-partitioned into Tau condensates (Figure 4A). However, RNA-containing condensates (Tau:RNA and Tau:PEG:RNA) were rich in either polyA or tubulin, and Tau:PEG:RNA condensates had subdomains rich in either polyA or tubulin. The average relative concentration of RNA (normalized to Tau on an individual condensate level) in Tau:RNA and Tau:PEG:RNA condensates significantly decreased upon addition of tubulin, whereas the average concentration of tubulin was higher in condensates containing RNA compared to no RNA (Figure 4B). Most smaller condensates were rich in tubulin, whereas the preformed large condensates had a higher RNA content (Figure 4C), which could result from the *de novo* formation of new smaller Tau:tubulin condensates. These data show a competition between RNA and tubulin for the co-condensation with Tau, whereby Tau can function as a homogenous ‘liquid matrix’ that hosts both competetive interaction partners in distinct subdomains (Figure 4A, bottom panel). A lower c_sat_ of Tau for the co-condensation with tubulin, compared to RNA, could explain the formation of new condensates that are rich in tubulin and low in RNA. In fact, tubulin appears to efficiently induce Tau LLPS through co-condensation, similar to coacervation observed in Tau:RNA systems (Figure 4A, top panel).

**Figure 4.**
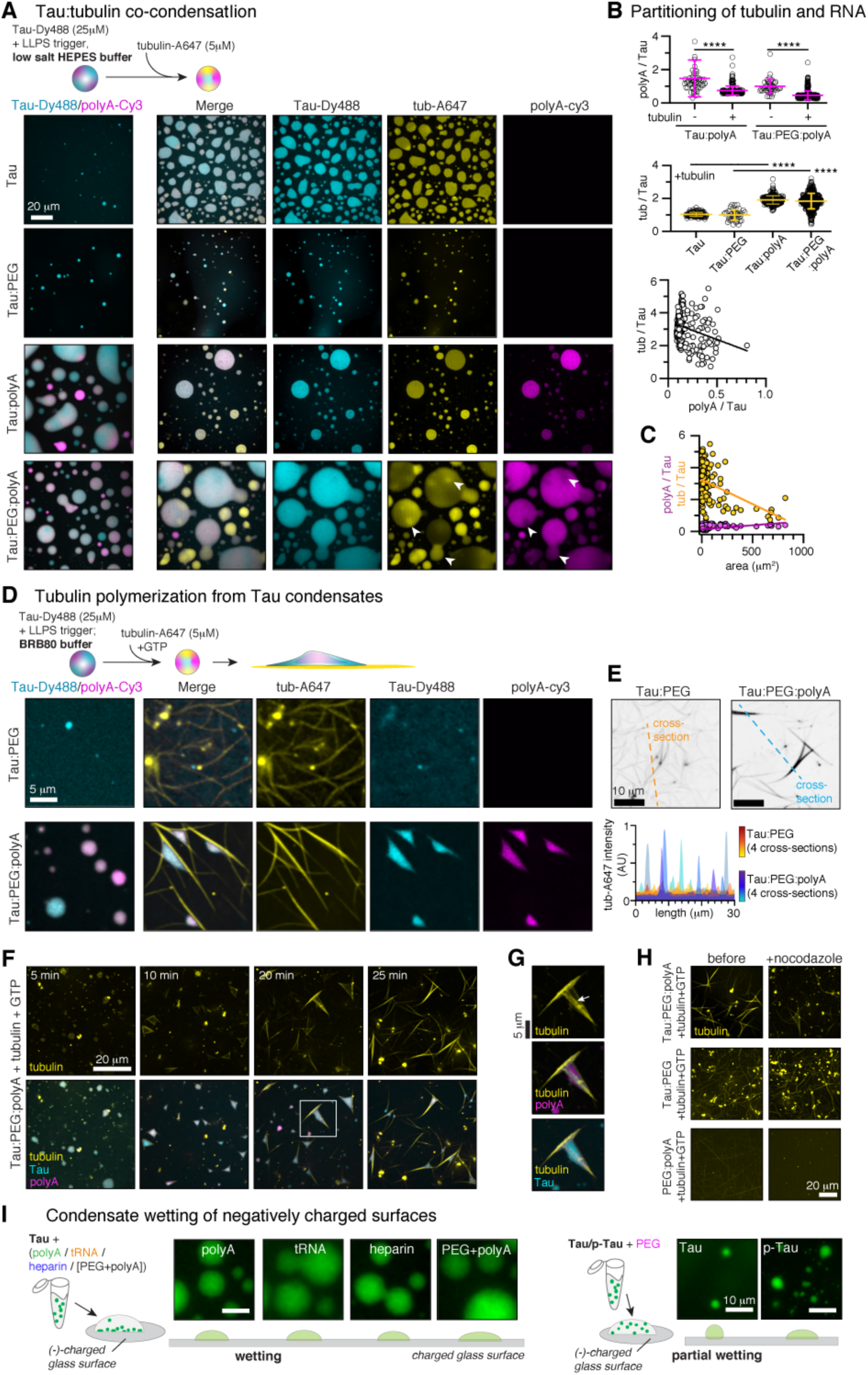
Tubulin and RNA compete for condensed Tau. A Tubulin co-condensation with Tau. Addition of tubulin-HiLyte647 (5 μM, 10:1 unlabeled:labeled tubulin) to a Tau-Dy488 solution (25 μM) induces Tau condensation. Addition of tubulin to Tau:PEG, Tau:RNA (polyA-Cy3), and Tau:PEG:RNA condensates leads to partitioning of tubulin into existing condensates and the occurrence of additional small Tau:tubulin condensates. Condensates with high tubulin-HiLyte647 intensity have low polyA-Cy3 intensity, and vice versa. In Tau:PEG:polyA condensates, individual condensates host RNA- and tubulin-rich domains (arrow heads) at homogeneous Tau-Dy488 intensity. Scale bars=20 μm. B Average intensity of RNA (top graph) normalized to Tau intensity in individual condensates. Addition of tubulin reduces the average amount of RNA in condensates. Intensity of tubulin (middle graph) normalized to Tau intensity in individual condensates. The presence of RNA increases the amount of tubulin in the condensates. Data shown as mean±SD. Data +/- tubulin and +/- polyA were compared by two-tailed Student’s t-tests. Tubulin and RNA intensity correlate negatively on an individual condensate level (bottom graph). Line represent linear regression to all data. C Condensate size (are on surface) relative to tubulin (yellow data points) and RNA (pink data points) intensity. Smaller (new) condensates are high in tubulin, larger (old) condensates have more RNA. Lines represent linear regressions to tubulin (yellow) and RNA (pink) data. D MT polymerization out of Tau:PEG and Tau:PEG:RNA condensates. Addition of tubulin-HiLyte647 (5 μM, 10:1 unlabeled:labeled tubulin) and GTP to Tau-Dy488 (25 μM) diluted in BRB80 polymerization buffer induces MT polymerization. Images taken 30 min after addition of tubulin show thin partially curved MT bundles in Tau:PEG system, and thick bundles attached to persisting Tau:RNA condensates in the Tau:PEG:RNA system. Scale bar=5 μm. E Arrangement (see MT bundles outlined with black lines) and tubulin-HiLyte647 fluorescent intensity (see line plots from tubulin channel) of MT bundles differ between Tau:PEG (top, yellow/red line plots) and Tau:PEG:RNA (bottom, blue/green line plots) systems. Four line plots at random positions are shown per condition. F Time course of MT polymerization out of Tau:PEG:RNA condensates. Scale bar=20 μm. G Close-up of Tau condensed phase connecting RNA condensate with MT bundles. Scale bar=5 μm. H Nocodazole treatment (120 μM for 30 min) of MTs formed from Tau:PEG:RNA and Tau:PEG condensates. MT formed in absence of Tau shown as control. Scale bar=20 μm. I Attachment of Tau condensates to negatively-charged (glutamate treated) glass surfaces. Tau:RNA and Tau:PEG:RNA condensates show pronounced surface wetting, whereas Tau:PEG condensates stay in solution or show partial wetting. p-Tau:PEG condensates also wet the surfaces. Scale bars=10 μm.

To investigate how the competition between RNA and tubulin affects MT polymerization from Tau condensates, we formed Tau:PEG and Tau:PEG:RNA condensates in MT polymerization buffer (BRB80) and then added tubulin together with GTP to enable tubulin polymerization. Tau:RNA coacervates did not form in BRB80 buffer due to their high sensitivity to the buffer composition (Expanded View Figure EV3A). After 30 min, MT assembly was observed in both Tau:PEG and Tau:PEG:RNA systems (Figure 4D). In the Tau:PEG system, many thin and in part curvy MT bundles formed on the surface of the imaging dish (Figure 4E) and a homogenous Tau-Dylight488 signal indicated no obvious colocalization of Tau with MTs at the resolution of the images taken (spinning disk confocal microscope, 60x objective). In Tau:PEG:RNA systems, however, we observed thick MT bundles with a unique spatial organization of bundles that were arranged to maximize the interaction surface with the remaining condensates containing Tau and RNA.

Neither Tau nor RNA signal was detected in the background, indicating that most Tau and RNA was condensed. Time course imaging revealed that the MT bundles formed out of Tau:RNA:tubulin condensates, initially aligned in a triangluar arrangement around the remaining Tau:RNA condensates, and then further grew in length and thickness (Figure 4F). Tau:RNA condensates remained attached to the MT bundle surface, and the condensed Tau phase seemed to ‘encapsulate’ both the RNA-containing condensates and MT bundle surface, thereby connecting these two structural entities (Figure 4G). The stability against nocodazole-induced disassembly was higher for Tau:PEG compared to Tau:PEG:RNA induced MT, possibly because free Tau could bind and stabilize MTs in Tau:PEG systems, whereas all Tau was condensed and not available for MT stabilization in Tau:PEG:RNA systems (Figure 4H). These data could explain early observations, in which a competition between tubulin and RNA for Tau was suggested based on the ability to isolate cellular Tau together with MTs only in the absence of RNA (Bryan *et al*, 1975).

Notably, condensates formed with p-Tau (from insect cells) in binary p-Tau:PEG systems remained intact upon addition of tubulin, attached to MTs, and showed MT surface wetting (Expanded View Figure EV3B; (Hernández-Vega *et al*, 2017). In tertiary p-Tau:PEG:polyA systems, however, p-Tau:RNA condensates showed similarly MT bundling and bundle coordination as observed using non-phosphorylated Tau in Tau:PEG:RNA systems.

The spreading of liquids on hydrophilic surfaces – the wetting behavior – is determined by the force balance between cohesive forces (F_C_) of molecules in the liquid phase that prevent wetting, versus attractive forces (F_A_) towards the surface that promote wetting (Chen & Bonaccurso, 2014). In this context, our data show that MT bundles arrange around Tau:PEG:RNA in a way that maximizes the interaction surface with the condensate while the droplet-shape of the condensates is minimally disturbed and only little spreading of condensates onto MT surfaces occurs. This indicates that cohesive forces in Tau:PEG:RNA condensates are stronger than the attractive forces towards MT surfaces, which are yet strong enough to guide spatial MT bundle orientation. In contrast, Tau:PEG condensates show no attachment to MTs, hence their attraction to the MT surface is minor. p-Tau:PEG condensates are wetting onto MT surfaces, hence have a high attraction to their surface at low cohesiveness.

We were able to confirm these ideas in a simplified model of Tau interactions with negatively charged surfaces - in analogy to the interaction of Tau with polyanionic MT surfaces. We prepared different Tau condensates and added them onto negatively charged (glutamate-treated) glass surfaces. Tau:PEG:RNA condensates, as well as binary Tau:RNA coacervates formed with different RNAs and heparin, rapidly attached to and showed liquid-like wetting of the negatively charged surfaces (Figure 4I). p-Tau:PEG condensates also spread onto charged glass. In contrast, Tau:PEG condensates showed little surface attachment with only partial wetting. Similar results were obtained for the interaction of condensates with positively charged (amine-treated) glass surfaces (Expanded View Figure EV3C).

These data indicated that Tau condensates containing negatively charged polymers – RNA or p-Tau – have a high affinity to charged surfaces hence a high (F_A_:F_C_)-ratio. PEG-induced Tau condensates show little wetting of charged surfaces, hence have less attractive and high cohesive forces and, thus, a low (F_A_:F_C_)-ratio. The wetting behavior of Tau condensates on charged surfaces may have important implications for their cellular functions, for example the binding to MTs (Tan *et al*, 2019; Siahaan *et al*, 2019; Mitchison, 2020; Hernández-Vega *et al*, 2017) or lipid membranes (Majewski *et al*, 2020; Mari *et al*, 2018; Snead & Gladfelter, 2019), but also for their capability to bundle and spatially organize MTs in the cytosol.

### Crowding, but not polyanions, induce gelation of Tau condensates

In addition to functional MT binding and bundling, Tau condensates have also been implicated in the generation of pathological Tau species that have the ability to seed Tau aggregation (Wegmann *et al*, 2018; Boyko *et al*, 2020; Kanaan *et al*, 2020). Protein condensates can undergo a phase transition from liquid to gel-like (Jawerth *et al*, 2020), which is thought to be the underlying mechanism for the development of protein aggregates from condensates. This aging behavior can be envisioned as a polymerization process, which can be described as “percolation” by the Flory-Stockmayer theory (Flory, 1941; Stockmayer, 1944; Choi *et al*, 2020b). Liquid-to-gel transitions were previously suggested for condensates of the RNA binding proteins FUS (Patel *et al*, 2015) and hnRNPA1 (Molliex *et al*, 2015), and for Tau in the presence of molecular crowding by us and others (Wegmann *et al*, 2018; Kanaan *et al*, 2020). Whether Tau coacervates show a similar transition in their polymer state is not known.

Percolation is accompanied by a decrease in molecular diffusion that can be measured as an age-dependent decrease in FRAP in condensates containing fluorescently labeled proteins. In Tau condensates (∼1% fluorescently labeled Tau-Dylight488) formed in Tau:PEG and Tau:PEG:RNA systems, we found the previously reported decay of recovery rate and increase in immobile molecules with time, which suggests percolation of these condensates (Figure 5A, left and right panels). FRAP of differently sized ROIs within Tau:RNA condensates showed that the time to half maximum fluorescence recovery, *t_1/2_*, correlated with the ROI size (Expanded View Figure EV4A), indicating that the recovery was dominated by molecular diffusion of Tau-Dylight488 (McSwiggen *et al*, 2019). Fully bleached condensates showed less recovery than partially bleached ones, suggesting that the recovery is dominated by molecular diffusion of Tau-Dylight488 inside condensates. When recording FRAP at different time points (1h, 4h, 24h), we found that binary Tau:RNA coacervates (without PEG) showed vivid fluorescence recovery (≥60% mobile molecules) even after 24h, with only a small reduction in the recovery rate (Figure 5A, middle). This indicated that most Tau molecules maintained their diffusion, hence no percolation occurred in Tau:RNA condensates. Other binary Tau:polyanion coacervates (with tRNA and heparin) showed a similar maintained FRAP (Expanded View Figure EV4B).

**Figure 5.**
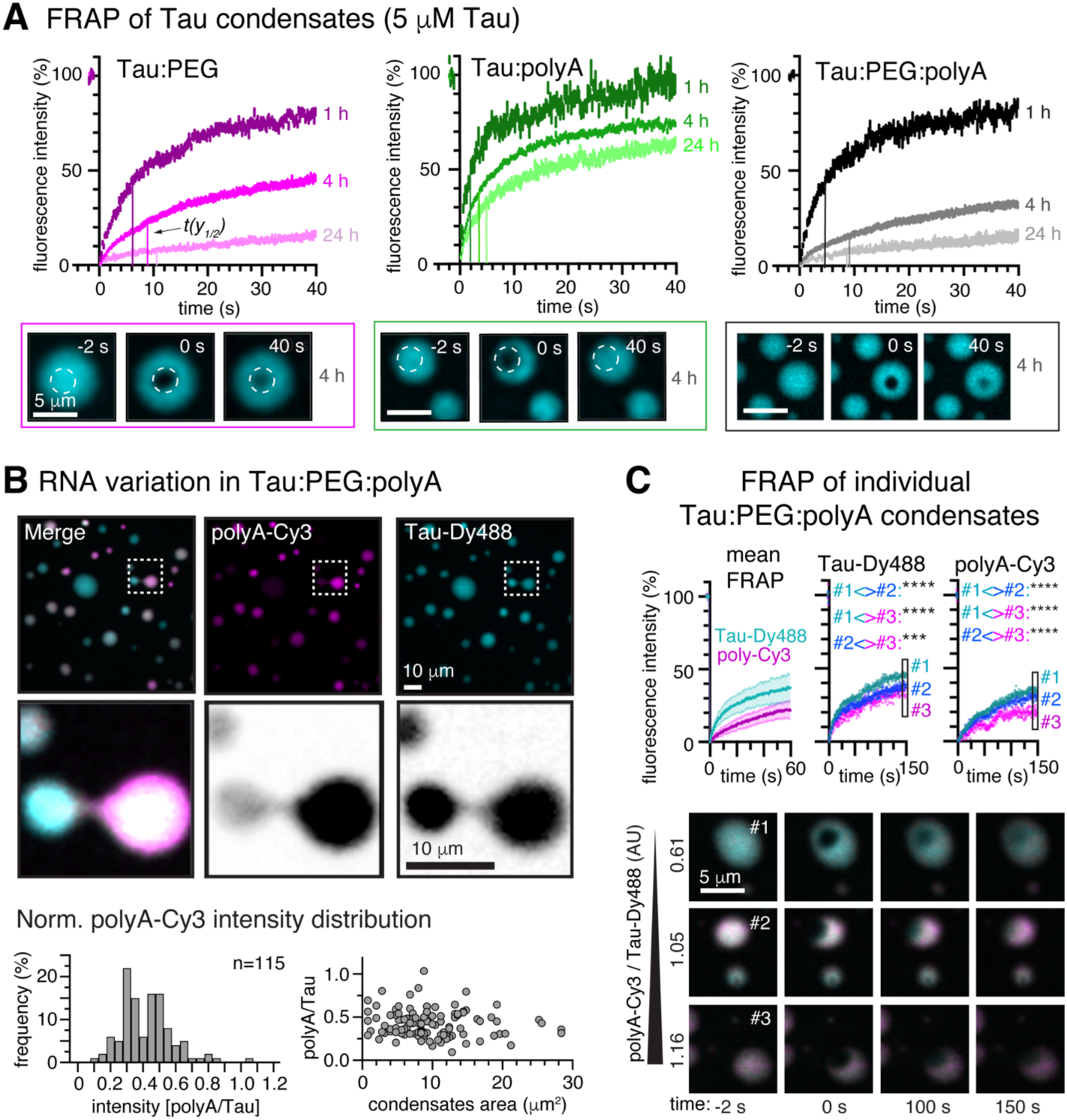
Tau coacervates only percolate in the presence of PEG. A FRAP of Tau:PEG, Tau:polyA, and Tau:PEG:polyA condensates at 1h, 4h, and 24h after LLPS induction. Tau:PEG and Tau:PEG:polyA condensates show a decrease in recovery rate and mobile fraction over time. Tau:polyA coacervates maintain fast recovery and a large mobile fraction even after 24h. Time until half maximum recovery: t(y_1/2_)_polyA_: 1h≈2s, 4h≈3.5s, 24h≈4.5s; t(y_1/2_)_PEG_: 1h≈6s, 4h≈8s, 24h≈11s; t(y_1/2_)_PEG+polyA_: 1h≈4.5s, 4h≈9.5s, 24h≈9s). Scale bars=5 μm for Tau:polyA and Tau:PEG, and 10 μm for Tau:PEG:polyA. Data presented as mean±SEM, n=15 to 20 condensates per condition and time point. B Microscopy of Tau:PEG:polyA condensates (Tau-Dylight488 and polyA-Cy3) shows condensates with different Cy3 intensities. The bimodal fluorescence intensity distribution suggests two groups of condensates with high or low Cy3 fluorescence. Cy3-intensities, normalized to Tau-DyLight488 intensities in the same condensates, were independent of condensate size. Scale bar=10 μm. C FRAP of Tau-Dylight488 and polyA-Cy3 of 4h-old Tau:PEG:polyA condensates. The mean FRAP of polyA-Cy3 is lower than of Tau-Dy488 (left). Data shown as mean±SEM, n=24 condensates. FRAP data of individual condensates (bottom; #1 to #3) shows a decrease of Tau=Dy488 (middle) and polyA-Cy3 (right) FRAP with an increase of polyA-Cy3 (relative to Tau-Dy488) in the condensates. Scale bar=5μm.

Interestingly, using fluorescently labeled Tau (Tau-Dylight488) and polyA (polyA-Cy3), we observed condensates with different Cy3-fluorescence intensities in Tau:PEG:RNA coacervates (Figure 5B), whereby the Cy3-intensity did not correlate with condensate size. Performing FRAP of Tau and RNA in 4h-old condensates, we found an overall low recovery of Cy3-RNA (Figure 5C, Expanded View Figure EV3C), but FRAP of individual condensates suggested some negative correlation between the relative RNA fluorescence (ratio Cy3-polyA:Tau-Dy488) and fluorescence recovery of Tau-Dy488 and polyA-Cy3.

The condensed (high density) and the light (low density) phase in Tau condensation samples can be separated by centrifugation (1.6×10^3^g for 15 min at room temperature). We noticed that binary Tau:polyA coacervates, and coacervates formed with other polyanions (heparin, tRNA, polyU), did not separate from the rest of the solution (Expanded View Figure EV4D), whereas Tau:PEG and Tau:PEG:RNA condensates could be pelleted under the same condition. This suggested that binary Tau:polyanion coacervates have a lower density compared to PEG-induced condensates.

In summary, Tau:RNA coacervates appear to have a density similar to aqueous solutions and maintain internal fast molecular diffusion over time, whereas crowding by PEG induces percolating condensates of higher density. In the presence of polyanions and crowding, Tau condensates with different polymer properties can co-evolve. We envision that the ensemble of different coexisting Tau coacervates will further increase with the number of Tau interactions partners in the cell.

### Decoupling of Tau seeding potential and percolation in Tau coacervates

PEG-induced Tau condensation can induce the formation of oligomeric seeding-competent Tau species (Wegmann *et al*, 2018; Kanaan *et al*, 2020). Since percolation is thought to be a prerequisite for the formation of aggregates in protein condensates, we hypothesized that binary Tau:RNA coacervates would be less efficient in producing such oligomeric Tau species, and therefore also less efficient in initiating Tau aggregation. To test this idea, we produced Tau:PEG, Tau:RNA (tRNA), and Tau:PEG:RNA condensates and examined the formation of SDS-stable higher molecular weight (HMW) Tau species (e.g. dimers, oligomers) over time by SDS-PAGE (Figure 6A). High concentrations of heparin (287μg/ml; hep_AGG_) induce β-structured Tau aggregates (Goedert *et al*, 1996; Kampers *et al*, 1996), which were included as positive controls for HMW Tau formation. After 24h, all LLPS conditions, but not Tau alone, contained SDS-stable dimers and higher molecular weight (HMW) Tau similar to hep_AGG_. After 72h, all samples, including Tau alone, contained SDS-stable dimers and oligomers. The formation of HMW Tau species was further confirmed by SemiNative PAGE (Expanded View Figure EV4E). Lower concentrations of Tau (5 μM) also produced SDS-stable dimers in all LLPS conditions after 24h (Expanded View Figure EV4F). The formation of Tau oligomers in 24h-old Tau coacervates could be confirmed by dot blot analysis using antibodies with preferential binding to Tau oligomers (antibody 2B10 (Chandupatla *et al*, 2020); tRNA_LLPS_: p=0.0403 compared to Tau only; Expanded View Figure EV4G). These results show that Tau:RNA coacervates can produce SDS-stable Tau dimers and oligomers with similar or even higher efficiency as PEG-induced condensates. Comparable results were obtained for condensates prepared with the FTD-mutant pro-aggregant Tau^ΔK280^ (Expanded View Figure EV5A-D).

**Figure 6.**
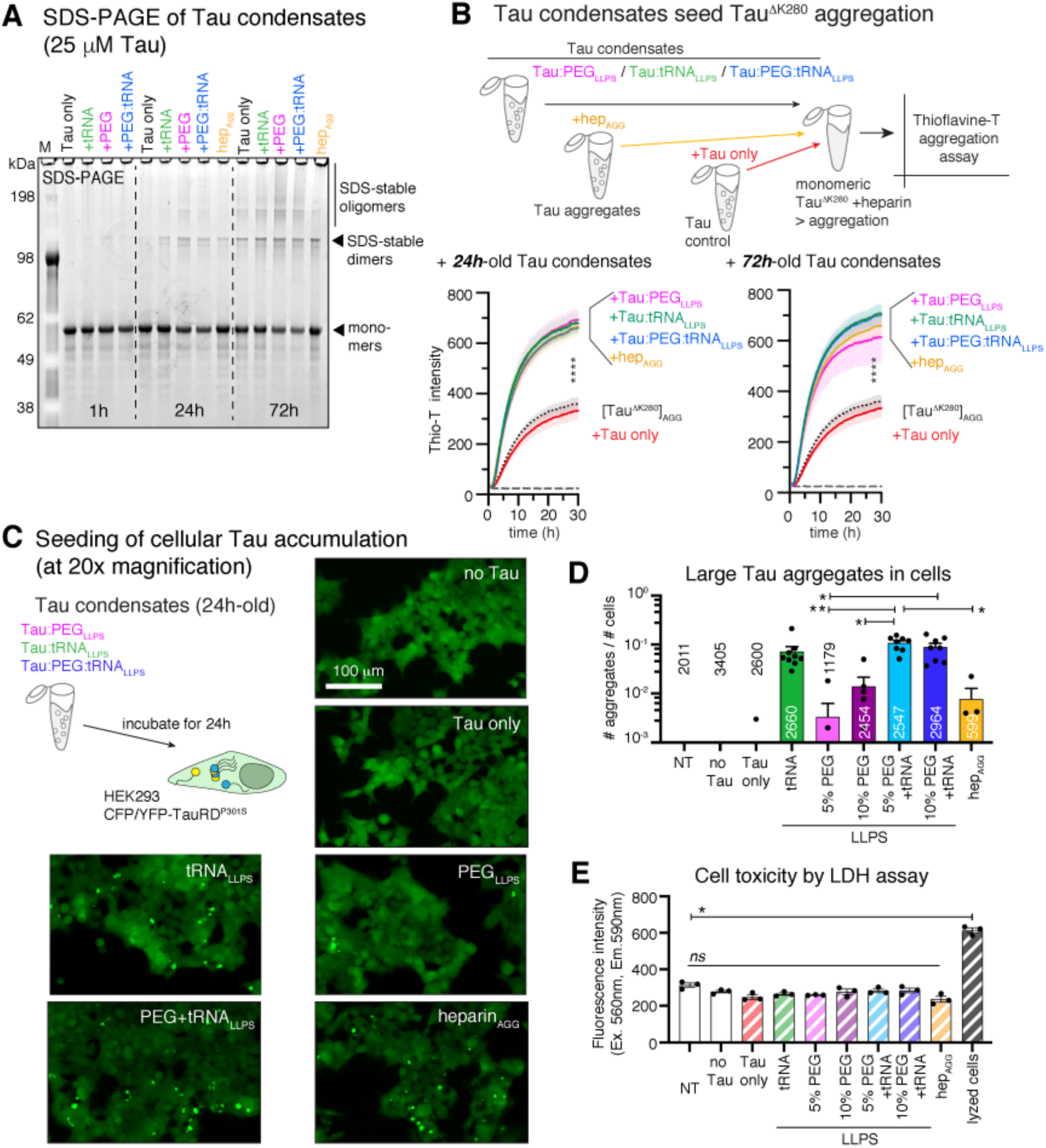
Tau coacervates and condensates produce Tau species with seeding potential. A SDS-PAGE of Tau incubated in LLPS (PEG, tRNA_LLPS_, PEG:tRNA_LLPS_) or aggregation (hep_AGG_) conditions for 1h, 24h, or 72h. SDS-stable Tau dimers and oligomers species evolve in LLPS and hep_AGG_ samples after 24h. After 72h, all conditions contain SDS-stable Tau oligomers and dimers. B Thioflavin-T assay for the enhancement of *in vitro* Tau aggregation by aged Tau condensates. Tau^ΔK280^ (10 µM) incubated with heparin (molar ratio Tau:heparin=4:1; [Tau^ΔK280^]_AGG_) was used as base line condition for Tau aggregation. Addition of 24h-old or 72h-old Tau LLPS samples (9.2 µg/ml=0.2µM Tau) produced an increase in Tau^ΔK280^ aggregation for all LLPS conditions (PEG, tRNA, PEG:tRNA) and hep_AGG_. Addition of Tau only did not affect Tau^ΔK280^ aggregation. Data shown as mean±SD, n=3. One-way ANOVA with Tukey test for multiple comparison for values at 30h. C HEK sensor cell Tau aggregation assay. Cells were treated with 2h-old condensates or aggregates (1.2 µM Tau in the presence of 0.8% lipofectamine). The number of Tau aggregates was determined from low magnification (20x) images taken in the green channel and normalized to the number of cells (DAPI). Scale bar=100 μm. D Quantification of seeding assay in (C). Data were collected from three images of 3 wells for each condition. Data shown as mean±SD, n=3. One-way ANOVA with Tukey test for multiple comparison. E Cell toxicity (LDH assay) induced by treatment of HEK sensor cells with condensates or aggregates. Data shown as mean±SD. One-way ANOVA with Tukey test for multiple comparison.

The potential to induce misfolding and aggregation is an important parameter of the pathological activity of Tau species (Soto & Pritzkow, 2018). To assess whether Tau dimers/oligomers that evolved from Tau condensates can induce Tau aggregation, we monitored the reaction by Thioflavin-T where the fluorescence intensity correlates with the increase of β-structure (Figure 6B). Monitoring the aggregation of pro-aggregant Tau^ΔK280^ with heparin ([Tau^ΔK280^]_AGG_) in the presence or absence of small amounts (9.2 µg/ml≍0.2 µM Tau) of 24h-old or 72h-old Tau condensates, we found that binary Tau:RNA (Tau:tRNA), Tau:PEG (Tau:PEG, and Tau:PEG:RNA (Tau:PEG:tRNA condensates all significantly increased the aggregation of Tau^ΔK280^ to levels achieved by addition of pre-formed Tau aggregates (hep_AGG_). Seeding of Tau^ΔK280^ with condensates prepared from Tau^ΔK280^ gave similar results (Expanded View Figure EV5E).

To analyze if the *in vitro* seeding potential of Tau condensates would translate into seeding “bioactivity” in cells, we treated HEK sensor cells stably expressing pro-aggregant Tau molecules (Tau repeat domain with P301S mutation) fused to CFP and YFP (HEK293 CFP/YFP-TauRD^P301S^; (Holmes *et al*, 2014)) with 24h-old Tau condensates (1.2 μM Tau in culture medium including 1% lipofectamine for 2 h). In these cells, local Tau inclusions, which depend on the material used for seeding, can be triggered upon exposure to seeding-competent Tau material (Kaufman *et al*, 2016). To our surprise, Tau:RNA coacervates – although not percolating over time – induced Tau aggregation more efficiently than PEG-induced condensates and pre-aggregated Tau, and similar to Tau:PEG:RNA condensates (Figure 6C,D). Tau:heparin condensates had even higher seeding potential (Expanded View Figure EV6A). The cell toxicity was similar across all condensate types and aggregates (Figure 6E). 1h-old condensates did not induce Tau aggregation in HEK sensor cells (Expanded View Figure EV6A).

These data show that, in a day or maybe even faster, different condensed Tau phases can evolve into Tau species that contain structural or biochemical information sufficient to seed *in vitro* and *in cellulo* Tau inclusions. Importantly, the templating Tau species can evolve without percolation of the condensates. Tau protein aging due to non-enzymatic fragmentation and amino acid racemization was shown to shape the composition of higher molecular weight Tau species (Watanabe *et al*, 2004), and similar processes may also contribute to the formation of seeding-competent Tau in aged Tau condensates.

### Tau condensates induce Tau accumulation around the cell nucleus

Tau oligomers and aggregates (Kaufman *et al*, 2016), and even monomers (Sharma *et al*, 2018), can seed distinct patters of Tau inclusions (“strains”), which are encoded in the nature (conformation and composition) of the seeding Tau species. We assessed which types of Tau accumulations would be seeded by Tau condensates and compared them to Tau aggregate induced Tau accumulations.

Treatment of HEK sensor cells with condensates resulted in numerous Tau accumulations, which we categorized into three types according to their cellular localization - with respect to the nucleus - their fluorescent intensity, and their size (Figure 7A-D; Expanded View Figure EV6B). Notably, multiple accumulations of the same or different type could occur in the same cell. Large, bright cytoplasmic inclusions (CYT Tau) located adjacent to the nucleus in the cytoplasm (∼1.5 μM distance to nuclear envelope). Spherical, often bright accumulations residing inside the nucleus (NUC Tau) in areas of low chromatin density (=propidium iodide-negative areas). And small, dimmer Tau accumulations at the nuclear envelope (NE Tau) formed a thin non-uniform layer of small granule-like structures around the nucleus and colocalized with lamin and nuclear pore proteins in the nuclear envelope. CYT Tau and NUC Tau accumulations have previously been characterized in these cells (Holmes *et al*, 2014) and were also shown to also contain RNA (Lester *et al*, 2021). In contrast, NE Tau has not been reported yet, probably because of its 4-fold lower fluorescence (Figure 7B) and its ‘invisibility’ to amyloid dyes (Figure 7E).

**Figure 7.**
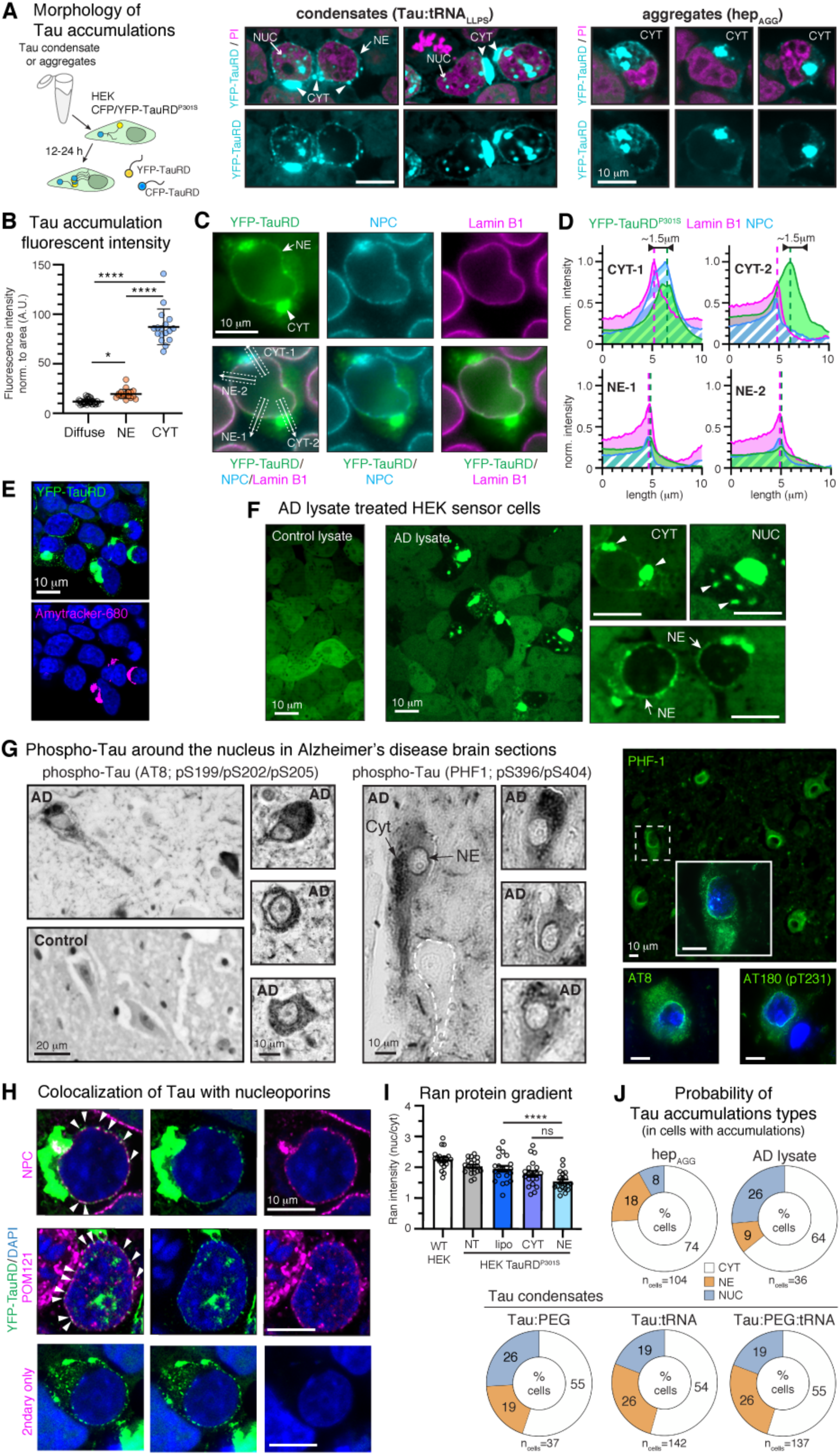
Nuclear envelope Tau accumulations seeded by condensates and in AD brain. A Confocal microscopy of HEK sensor cells treated with 24h-old Tau condensates (Tau:tRNA) shows different types of intracellular CFP/YFP-Tau accumulations: spherical intranuclear Tau (NUC), bright cytoplasmic accumulations (CYT) adjacent to the nucleus, and small, dimmer accumulations right at and along the nuclear envelope (NE). Cells treated with Tau aggregates (hep_AGG_) had mainly large cytoplasmic (CYT) Tau inclusions. The different Tau accumulations coexisted in most cells, and also more than one accumulation of the same type could occur. Nuclei were stained with propidium iodide (PI, pink). Scale bars=10 μm. B Fluorescence intensity of NE and CYT Tau compared to soluble Tau in the cytosol of cells without accumulations. Data shown as mean±SD. One-way ANOVA with Tukey test for multiple comparison. C Immunolabeling of lamin-B1 and nuclear pore complexes (NPC) shows the location of CYT and NE Tau in relation to the nuclear envelope in seeded HEK sensor cells. White arrows and boxes show position and width of cross-sectional profiles shown in (D). Scale bar=10 μm. D Cross sectional profiles show the localization of Tau (green) in CYT accumulations adjacent to the nucleus in the cytosol, and the colocalization of NE Tau with lamin (pink) and NPCs (blue). Notably, CYT-1 also contains NPC signal. E Seeded HEK sensor cells stained with the amyloid dye (Amytracker-680). Scale bar=10 μm. F HEK sensor cells treated with AD brain TBS extract show similar CYT, NUC, and NE accumulations as Tau condensate treated cells. Scale bars=10 μm. G Immunolabeling of phosphorylated Tau (PHF1=pS396/404, AT8=pS199/pS202/pT205, AT180=pT231) in AD brain section. Both 3,3′-Diaminobenzidine (DAB) and fluorescence labeling show Tau accumulating in a layer at the nuclear envelope (NE) in non-tangle neurons with or without cytosolic (CYT) Tau accumulations. Control brain does not show Tau labeling in neuronal cell bodies. Scale bars=10 μm. H High resolution confocal images show colocalization (arrow heads) of small NE Tau clusters with nuclear pore proteins (NPC marker, POM121). Scale bars=10 μm. I Reduced nucleo-cytoplasmic Ran gradient (nuclear/cytosolic Ran intensity) in HEK sensor cells having CYT Tau (CYT), or NE Tau (NE), compared to non-treated (NT) or control treated (lipo) sensor cells. Wildtype (WT) HEK cell Ran gradient was analyzed for the effect of TauRD^P301S^ expression in general. Data shown as mean±SD. One-way ANOVA with Tukey test for multiple comparison. J Frequency of Tau accumulation types observed in sensor cells treated with Tau aggregates (hep_AGG_), AD lysate, or Tau condensates. In cells having Tau accumulations we determined which accumulation types were present, regardless of the co-presence of other accumulation types and if multiple accumulations of the same type were present. n=number of cells with accumulations analyzed, resulted from the actual number of cells in the three replicates.

To see if NE Tau would also have a relevance in the brain, we treated HEK sensor cells treated with human AD brain lysate and determined which Tau accumulations would develop. AD, but not control, lysate treated cells showed all three types of Tau accumulations (CYT, NUC, NE; Figure 7F). We observed similar types of Tau accumulations also in primary neurons expressing CFP/YFP-TauRD^P301S^ that were treated with AD lysate (Expanded View Figure EV6C). Immunostainings of brain sections - visualized with either DAB or fluorescence - confirmed that phospho-Tau (AT8, PHF-1, and AT180 epitopes) is accumulating around neuronal nuclei also in AD brains (Figure 7G). It was suggested that Tau at the NE interacts with nuclear pore complexes, thereby impairing the nucleocytoplasmic transport (NCT) of proteins (Eftekharzadeh *et al*, 2018) and RNA (Cornelison *et al*, 2019). Similarly, in HEK sensor cells, NE Tau colocalized with nuclear pore proteins (Figure 7F), and, compared to cells having only CYT Tau, cells having NE Tau showed an even stronger decrease in the nuclear:cytoplasmic Ran protein gradient, a measure for nucleocytoplasmic transport deficits (Figure 7G).

Interestingly, we observed that HEK sensor cells treated with different Tau seeding agents – condensates, aggregates, or AD lysate – showed different amounts of the three Tau accumulations types. Whereas pre-formed fibrillary Tau aggregates (hep_AGG_) induced mostly CYT Tau accumulations, Tau condensates more frequently induced NE Tau and NUC Tau (Figure 7A+J; Expanded View Figure EV6B). AD lysate treated cells showed similar amounts of CYT and NUC Tau, but less NE Tau compared to condensate seeded cells.

In summary, these data suggest that aged Tau condensates contain Tau species that template different types of Tau accumulation in cells, similar to Tau from AD brain. The seeded Tau accumulations include Tau assemblies associated with the nuclear envelope that could interact with nucleoporins and inflict nucleocytoplasmic transport deficits. Since the emerging Tau accumulations in the used cell model depend on the nature of Tau in the seeding material (Kaufman *et al*, 2016), Tau condensates may infer distinct molecular conformations specifically templating NE Tau.

### Nuclear envelope Tau is loosely packed compared to cytosolic Tau aggregates

Tau accumulations at the nuclear envelope in HEK sensor cells did not stain with amyloid dyes (Figure 7E), and until now they were also not reported in numerous Thioflavin staining experiments of AD brains. We suspected that NE Tau may be less densely packed, i.e. not aggregated, and maybe even be in a liquid-condensed state.

A characteristic that distinguishes liquid-like biomolecular protein condensates from tightly packed aggregates is the molecular diffusion in their interior, and the exchange of molecules with the surrounding cytosol, which can be assessed through FRAP. We found that full bleaching of NUC Tau accumulations barely showed FRAP (Figure 8A,B), as would be expected for Tau aggregates, however, which could also result from the limited pool of soluble CFP/YFP-TauRD available for replacing bleached molecules. Full bleach of NE Tau accumulations did lead to a recovered up to ∼20%, indicating that Tau in these accumulations was mostly immobile (80%), however could to some extent be replaced by non-bleached CFP/YFP-TauRD molecules from the surrounding cytosol. Similar results were obtained upon full bleach of 4h-old Tau:PEG:RNA condensates (Expanded View Figure EV4A). Notably, NE granules were mostly immobile and without obvious fusion events (Expanded View movie M1), as would be expected for gel-like condensed Tau phases in the cytosol (Wegmann *et al*, 2018). Partial bleach of CYT Tau also lead to only 20% recovery, indicating some replacement of bleached molecules by non-bleached ones from the cytosol or through molecular diffusion inside accumulations. Of note, the recovery rate was much faster for NE than for CYT Tau inclusions (t_1/2_ for NE =2.5; t_1/2_ for CYT: t_1/2_=7 sec), suggesting a condensed phase behavior of NE Tau versus an aggregate behavior of CYT Tau.

**Figure 8.**
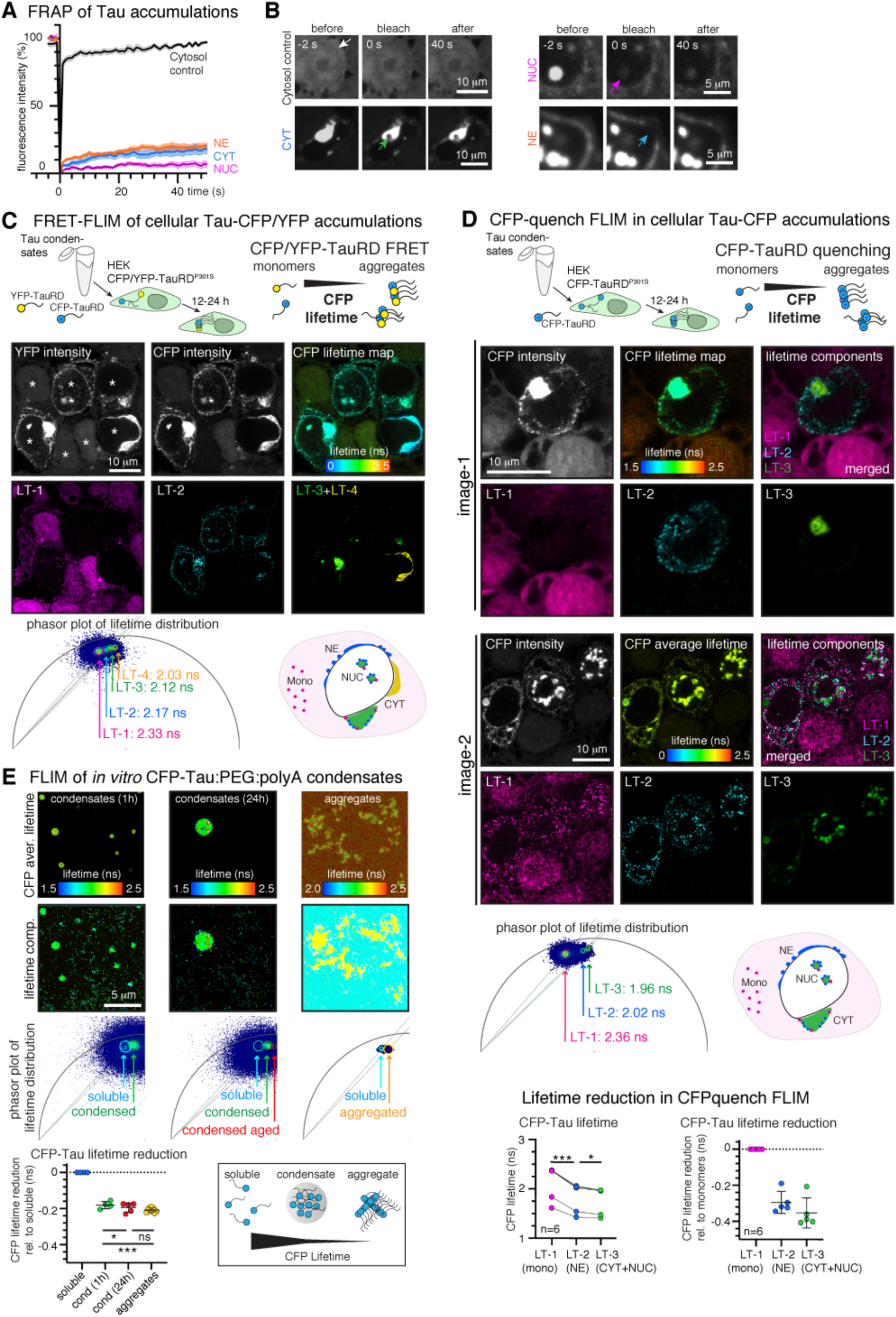
FRAP and FLIM reveal differences in molecular packing in Tau accumulations. A FRAP of Tau accumulations show no recovery of NUC Tau (NUC: <10% recovery; t_1/2_ ≍11 s), ∼20% recovery of CYT and NE Tau with a faster recovery rate for NE Tau (NE: t_1/2_ ≍2.5 s; CYT: t_1/2_ ≍7 s). Data shown as mean±SD, n= 15-20 cells per treatment group. B Example FRAP image series for cytosol and the different Tau accumulation types in HEK sensor cells treated with AD brain lysate (pre-bleach=-2s, immediately after bleaching=0s, after recovery=40s). Arrows indicate position of bleached ROIs. Scale bars=5 or 10 μm as indicated. C High-resolution CFP FLIM of HEK sensor cells expressing CFP/YFP-TauRD^P301L^ and treated with Tau condensates. FRET between CFP and YFP induces CFP-lifetime reduction in densely packed CFP/YFP- TauRD^P301L^ accumulations. Fit-free analysis of the lifetime distribution in phasor plots enables the assignment of distinct lifetime clusters (circles in phasor plot) to different Tau accumulation types. LT-2=NE Tau, LT-3=CYT and NUC Tau, LT-4=fully aggregated CYT Tau. LT-1 corresponds to soluble CFP- TauRD in the cytosol of cells without accumulations. Scale bars=10 μm. D CFP-FLIM of seeded HEK sensor cells expressing only CFP-TauRD^P301L^. The dense packing of CFP- TauRD in Tau accumulations induces fluorophore quenching that can be detected as decrease in CFP-lifetime. Fit-free analysis of lifetime phasor plots enables the assignment of distinct lifetime clusters (circles in phasor plot) to different Tau accumulation types. LT-2=NE Tau, LT-3=CYT and NUC Tau, and LT-1=soluble CFP-TauRD in the cytosol of cells without accumulations. Graphs show lifetimes of CFP-lifetime components assigned to soluble, NE, and CYT+NUC Tau in 6 different images (left graph, RM one-way ANOVA with Tukey post-test) and lifetime reduction relative to LT-1 in each respective image (right graph, one-way ANOVA with Tukey post-test.). Scale bars=10 μm. E FLIM of 1h- and 24h-old CFP-Tau (full-length) condensates and heparin induced CFP-Tau aggregates. Lifetime components are indicated in phasor plots. Graph shows lifetime reduction in condensates and aggregates relative to respective lifetime of soluble CFP-Tau. One-way ANOVA with Tukey post-test. Scale bar=5 μm.

**Figure 9.**
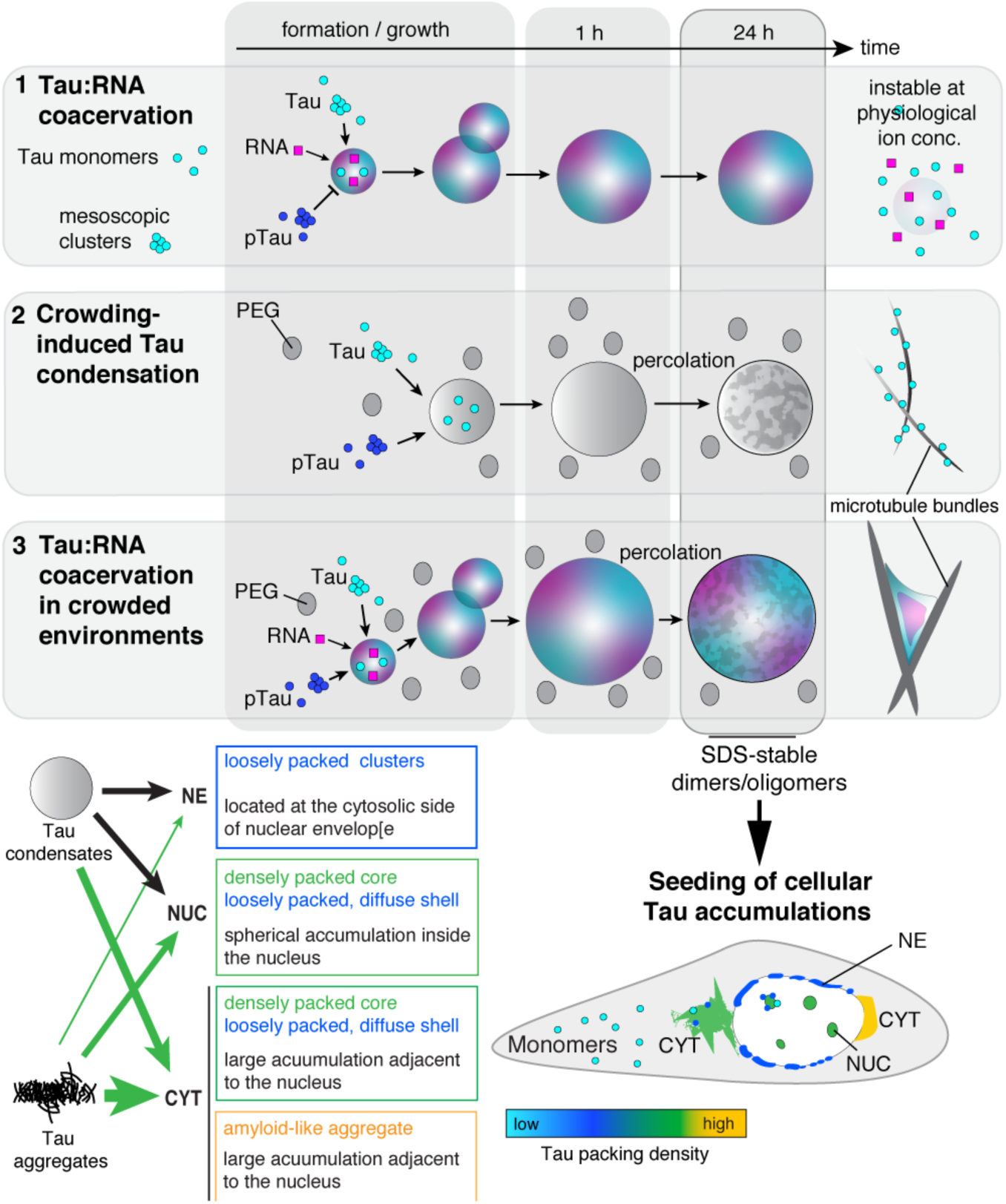
Tau condensation in different environments and condensate-induced cellular Tau accumulations. Tau coacervation with RNA (1) - or other polyanions – without PEG produces non-percolating condensates that are instable at physiological ion concentrations. Phosphorylation (pTau) inhibits coacervation of Tau. Crowding-induced condensates (2) are stable at physiological salt concentrations and percolate into gel-like condensates within 24 h. Tau:PEG condensates promote MT growth but disassemble during the polymerization process. Coacervation of Tau with RNA in the presence of molecular crowding (3), a condition that likely exists in the cytosol, produces condensates that are stable at physiological ion concentrations and also percolate into gel-like condensates. Tau:PEG:RNA condensates strongly promote MT bundling, and they persist as condensed phases attached to MT bundles, thereby inducing a distinct spatial organizing of MT bundles. Tau:PEG:RNA condensates can function as ‘liquid matrix’ that can host different otherwise not compatible Tau binding partners, such as RNA and tubulin. Regardless of their physiochemistry and origin, all condensates induce the formation of SDS-stable Tau dimers and oligomers with time, and efficiently induce Tau aggregation in sensor cell and neuron models. Tau condensates induce spherical accumulations in the nucleus (NUC), large aggregates in the cytosol (CYT) adjacent to the nucleus, and small granular Tau condensates at the nuclear envelope (NE), which in part colocalize with nucleoporins and impair the nucleocytoplasmic transport. In contrast, heparin-induced Tau aggregates mostly seed CYT accumulations.

To confirm our impression that NE Tau may resemble condensed Tau phases, we further investigated the molecular packing density of Tau in the different accumulations types used high-resolution CFP fluorescence lifetime imaging (FLIM). The decrease in fluorescent lifetime of the donor fluorophore in a FRET system, i.e. CFP-YFP, can be used as a measure for FRET-activity, which indicates the proximity (≤10 nm) between donor and acceptor fluorophores (Gorlovoy *et al*, 2009). In protein assemblies, the reduction in donor lifetime thus should correlate with the molecular packing density in the assembly.

FRET-FLIM imaging of HEK sensor cells expressing CFP/YFP-TauRD^P301S^ revealed reduced CFP lifetimes in all Tau accumulations compared to cytosolic soluble CFP-TauRD (Figure 8C). A fit-free analysis of the CFP-lifetime distribution in phasor plots enables a pixel-by-pixel assignment of lifetimes, regardless of the number of lifetime components relevant for the contained fluorescent molecules. Using this approach, we found distinct phasor plot lifetime clusters for CFP-TauRD in the cytosol (LT-1∼2.3 ns), in NE accumulations (LT-2∼2.2 ns), and in CYT and NUC Tau (LT- 3∼2.1 ns); a subset of CYT Tau had even lower CFP-lifetimes (LT-4∼2.0), probably resembling the fully aggregated Tau were also detecting by amyloid-dyes. The minor overall differences in CFP lifetimes between the different Tau accumulations indicated similar FRET efficiencies and suggest that the proximity of CFP and YFP in all types of accumulations may be ≤10 nm.

In addition, the dense packing of CFP-TauRD in the accumulations may lead to self-quenching of CFP, which would reduce the CFP lifetime independent of FRET with YFP acceptor molecules. In fact, it was shown that lifetime reduction of fluorescent (protein) tags to some extend correlates with the aggregation state of proteins such as TauRD or polyQ (Chen *et al*, 2017). Using this approach, we performed FLIM of HEK sensor cells only expressing CFP-TauRD but not YFP-TauRD (Figure 8D). Tau accumulations in these cells could be clearly distinguished from cytosolic Tau according to their CFP-lifetimes, indicating quenching of CFP. The fluorescent lifetime of CFP-TauRD in NE Tau (LT-2; reduction of -0.30 ns compared to soluble Tau) was significantly reduced compared to soluble cytosolic CFP-TauRD (LT-1), and CFP-TauRD in CYT and NUC Tau (LT-3) had an even stronger lifetime reduction (reduction of -0.35 ns compared to soluble Tau). Some CFP-Tau at the edge of dense CYT and NUC Tau had lifetimes similar to NE Tau, suggesting that these accumulations were surrounded by less densely packed Tau in a state similar to NE Tau. These data indicated that the packing density was lower in NE Tau than in CYT and NUC Tau, which would support the idea of NE Tau being in a liquid-condensed state.

To test whether the CFP-lifetime reduction in NE Tau would be similar in Tau condensates, we performed CFP-quench FLIM on *in vitro* CFP-Tau condensates (CFP-Tau:PEG:RNA; Figure 8E). In 1h-old condensate preparations, we found a reduced fluorescent lifetime of CFP-Tau in condensates compared to soluble CFP-Tau in the light phase (reduction of 0.19 ns compared). In aged condensates (24h-old), the lifetime in the condensed phase decreased a little further and reached a lifetime reduction (compared to the respective soluble CFP-Tau values in each preparation) similar to aggregated CFP-Tau fibrils prepared with heparin. Aging CFP-Tau condensates thus show a similar progressive CFP-lifetime reduction as observed between NE and CYT Tau accumulations in cells, suggesting that Tau at the nuclear envelope may indeed have a molecular packing like Tau condensates.

In summary, we find that CYT and NUC Tau accumulations are characterized by low FRAP and high FRET and CFP-lifetime quenching, which indicates low mobility and dense packing of molecules in their interior, reminiscent of aggregated Tau. In contrast, NE Tau shows faster FRAP and less FRET and CFP quenching, similar to condensates formed in the presence of molecular crowding and RNA *in vitro*. It is therefore possible that small granular Tau accumulations at the nuclear envelope may be in a liquid-condensed state.

## Discussion

In recent years, biomolecular condensation of Tau has mostly been studied by looking at isolated aspects of the phenomenon (MT binding versus pathological aggregation) and in very specific Tau LLPS systems – either coacervation with polyanions or crowding-induced condensation. Critical investigations questioning the existence of Tau condensates in the cellular environment are missing though, and two studies so far addressed the question of Tau condensation in cells (Zhang *et al*, 2020; Wegmann *et al*, 2018). To close this gap, we described the process of Tau condensation in LLPS systems with physiological relevance *in vitro*, and link these data to functional Tau condensation in the context of MT binding and polymerization, as well as to condensate-induced pathological Tau aggregation in cells.

To achieve this, we adapted different light-based techniques to study Tau condensation *in vitro* and in cells. The use of trDLS allowed to monitor the modulation of Tau condensation by ion composition, PTMs, and other biomolecules, i.e. RNA. With high-resolution FLIM we could image the molecular density, hence packing, of different types of Tau assemblies, and thereby created the possibility to distinguish Tau accumulations in cells based on their fluorescence lifetime characteristics. A simplified model of Tau condensates binding to charged glass surfaces allows predictions for the binding of Tau condensates to MTs (and maybe other cellular structures). Using these approaches in combination with techniques commonly used to study the condensation of proteins, such as turbidimetry, microscopy, and FRAP, revealed multiple new aspects of Tau condensation and its cellular functions.

### The importance of molecular crowding for Tau coacervation

We found that polyanions and molecular crowding drive the formation of liquid-like dense Tau phases with different formation kinetics, physicochemical properties, and interactions. At cytoplasmic ion concentrations, however, molecular crowding is essential to enable condensation of Tau and phospho-Tau. These findings provide a framework for Tau condensation in cytoplasm-like conditions, in which the interplay between molecular crowding, polyanions, and Tau PTMs needs to be considered.

Phospho-Tau, due to the additional negative charges of the phosphate groups, is unable to coacervate with RNA (at 0.005, 0.05, and 0.5 mg/ml polyA with 5 μM p-Tau) in the absence of crowding. Tau phosphorylation in cells is a complex and not well understood aspect of Tau (patho)biology that offers a fine-tuned response to diverse extracellular stimuli and intracellular signalling pathways. Although the contribution of phosphorylation to Tau LLPS remains mostly unclear, we gained evidence that some phospho-sites may act as pro-LLPS PTMs (i.e. pS262), whereas others may have little to no effect or even inhibit Tau condensation (i.e. pT205, pS404). Notably, other Tau PTMs can also tune Tau coacervation, such as acetylation (Ferreon *et al*, 2018; Ukmar-Godec *et al*, 2019), which further increases the complexity of Tau LLPS regulation in cells. In fact, Tau in cells is constitutively in a high state of phosphorylation, which makes it difficult to assign special functions to individual phosphorylation sites. Hence, specific and transient Tau phosphorylation in neuronal compartments, combined with local variance in RNA and crowding, could enable a context-dependent spatio-temporal regulation of neuronal Tau condensation. It would be interesting to test *in vitro*, for example, how stress-induced changes in the neuronal cytosplasm composition (e.g. Ca^2+^ and ROS increase, ATP decrease, Tau phosphorylation increase) influence Tau condensation in the presence of crowding, with or without RNA or other anionic Tau binding partners.

Notably, a colloidal pressure between 1-2 kPa and 20 kPa has been suggested for the cytoplasm (reviewed in (Mitchison, 2019)), which would correspond to the osmotic pressure exerted by ≤2.5% PEG8000 (Stanley & Strey, 2003). To robustly induce LLPS at physiological Tau concentrations (1-5 μM) we used 5% PEG8000 in most experiments. In such conditions, the size of PEG molecules (R_0_≍R_g_∼3.3 nm; (Soranno *et al*, 2014)) is about half that of the Tau monomers (R_0_=8-10 nm)(Mylonas 2008). Excluded volume effects exerted by PEG molecules encourage the interaction between Tau monomers, resulting in the formation of microscopic liquid condensates. Although similar effects could play a role for Tau assembly in the crowded cytosol, the interactions of Tau molecules with a number of cytosolic binding partners will substantially complicate this (simple) model in the cellular context.

### Tau condensates coordinate microtubules and other Tau binding partners

Tau coacervation with RNA is suggested to play a physiological role for the asociation of Tau with stress granules (Ash *et al*, 2021), and may also be involved in other RNA-associated cellular processes(Wheeler *et al*, 2019). Early efforts to assemble MTs from non-neuronal cells succeeded only when excess RNA (sequestering MAPs) was removed (Bryan *et al*, 1975). This suggested that RNA and tubulin compete for the binding to Tau, however, it reamined unclear how the competition would manifest. We show that RNA and tubulin seem to compete for the co-condensation with Tau, at least *in vitro*. The presence of RNA changes Tau-induced MT formation and the binding of Tau to MT bundles. Tau:PEG:RNA condensates are characterized by high cohesive forces that hold Tau and RNA molecules inside the condensed phase, yet they exhibit a strong affinity to negatively charged MT surfaces. The cohesivness lets Tau and RNA persist in a condensed phase during MT polymerization, while MT bundles that grow out of Tau:RNA condensates organize around the persisting condensates to maximize the condensate:MT contact area. The binding of Tau to other anionic cytoskeletal fibers and to the surfaces of lipid membranes (Mari *et al*, 2018; Majewski *et al*, 2020) may occur via Tau condensates as well. Interestingly, MT surfaces and RNA appear to occupy distinct and different volumes of the Tau condensates, suggesting that Tau condensates could function as a ‘liquid matrix’ that can host different binding partners that would otherwise not interact with each other. This could give Tau condensates an active role in coordinating and structuring cytoskeletal and other binding partners in the cytosol. For example, the pearl chain-like arrangement of chaperone accumulations along MTs could be enabled through a similar mechanism (Carrettiero *et al*, 2009).

The wetting of Tau:RNA condensates (with or without PEG) onto negatively charged MT surfaces can be described by fluid mechanics and may be interpreted as a form of coacervation, in which the polyanionic partner is not a soluble molecule but a stationary surface. To interpret the interactions of Tau condensates, one therefore needs to consider different interaction regimes and assembly state therories, i.e. single molecule interactions of Tau monomers with co-partitioning client molecules, as well as the polymer and liquid state interactions of Tau condensates. Notably, Tau condensates without RNA also induced MT formation but seemed to have little affinity for the MT surface; in this case MT assembly is probably supported by free soluble instead of condensed Tau, resulting in thinner MT bundles, a random MT organization and no remaining Tau condensates.

### Patho-physiological potential of Tau condensates in the absence of percolation

Biomolecular condensates of different neurodegenerative disease-associated proteins have been shown to progressively percolate and subsequently transit into aggregates (Ray *et al*, 2020; Patel *et al*, 2015; Molliex *et al*, 2015). We previously showed this path towards aggregation for crowding-induced p-Tau condensates (Wegmann *et al*, 2018). Our new data show an interesting deviation from this path for Tau coacervates. Binary Tau:RNA coacervates have a low density, show liquid-like surface wetting and coalescence for an extended time (multiple hours), and maintain fast molecular diffusion (seen by FRAP), even after a day. Hence, Tau coacervates did not behave like aging Maxwell fluids (Jawerth *et al*, 2020), a model recently suggested for condensates of FUS. Nevertheless, Tau:RNA coacervates developed SDS-stable Tau dimers and seeding-competent Tau species even more efficiently than percolating crowding-induced condensates. These findings show that, in Tau coacervates, the production of pathological Tau conformers is decoupled from condensate percolation. Aging Tau:RNA coacervates may thus need a classification different from RNA binding proteins. One possibility is that the SDS-stable dimers evolving in non-percolating Tau:RNA coacervates may function as small seeding Tau entities that condense but do not progressively establish interactions with other Tau molecules. Another explanation could be that Tau monomers can adopt a conformation specific for the condensed phase that is sufficient to gain Tau seeding activity. The idea of a distinct conformational state of Tau monomers encoding Tau seeding ability has previously been suggested (Mirbaha *et al*, 2018). However, we did not detect seeding potential in 1h-old coacervates, indicating that such monomeric conformation, if existent, would need more time to develop inside condensates.

All aged Tau condensates tested, despite their physicochemical differences, developed Tau seeding potential, indicating that intracellular liquid-condensed Tau phases need to be spatiotemporally well controlled to prevent their prolonged existence. Functional Tau LLPS may thus occur on short time scales (minutes to hours) and may be tightly regulated. For Tau, which has a lifetime of multiple days (Yamada *et al*, 2014) and binds to long-lived cellular structures, e.g. axonal MTs, lipid membranes, nucleoli, and nuclear pores, this seems a challenging task. Notably, the interaction of monomeric non-condensed Tau with MTs is very transient (Janning *et al*, 2014), Alternatively, the formation of Tau species with seeding potential could be prevented, for example, by molecules that partition into Tau condensates and exert chaperone function. How neurons balance functional Tau LLPS while preventing aberrant aging of Tau condensates needs to be further investigated.

### Condensed Tau at the nuclear envelope

In the human brain, as well as in cell models of Tau pathology, Tau accumulations can differ in their morphology, molecular packing, and localization. Cell models of seeded aggregation, such as HEK sensor cells, amplify even small seeding potential contained in samples from different sources. The accumulations evolving in these cells appear to depend on the templating Tau species and hence can have different characteristics. It is debated whether CFP/YFP-TauRD assemblies in sensor cells actually represent assembly forms found in human brain (Kaniyappan *et al*, 2020), because the fluorescent protein tag is thought to interfere with the assembly into amyloid-like fibrillar aggregates. Our data show that only a subset of large cytosolic aggregates in sensor cells adopt amyloid-like structure (stained with amyloid dyes) whereas the majority are other types of accumulations. However, also in brains of AD and tauopathy cases Tau accumulations other than amyloid-like paired helical filaments exist, i.e. Tau oligomer with high seeding activity.

Combining different microscopic techniques (confocal colocalization microscopy, FRAP, FRET-FLIM, fluorophore quenching FLIM) we generated evidence that Tau accumulating at the nuclear envelope may be in a condensed phase state as well. The ‘thin’ granular layer of Tau at the NE appears to be less densely packed compared to cytosolic and intra-nuclear Tau aggregates and exchanges Tau molecules with the cytoplasm, indicating a condensed liquid- or gel-like state of Tau. No fusion of NE Tau accumulations was observed, either because of time and spatial resolution limitations, or because progressed percolation prevented their fusion. We previously did not observe fusion of cytosolic Tau condensates in neurons either (Wegmann *et al*, 2018).

Using FRET-FLIM of CFP/YFP-TauRD sensor cells, and CFP-lifetime quenching FLIM of CFP-TauRD sensor cells, we show that all formed inclusions have a similar CFP-lifetime reduction because they all contain densely packed Tau molecules. Looking at mean FLIM or FRET does not allow to distinguish between accumulation types. However, using a fit-free analysis of the lifetime distribution we were able to clearly distinguish between types of accumulations. Liquid-condensed CFP-Tau has slightly less CFP-quenching than aggregated CFP-Tau due to the difference in molecular packing (dynamic molecules in condensates versus immobile, fixed molecules in aggregates). Amyloid-like aggregates show the highest CFP-quenching. Interestingly, we find that the quenching behavior in cells is similar to that of condensed CFP-Tau *in vitro* (Tau:PEG:polyA system), suggesting that this technique can be used to characterize the Tau assembling state in and outside of the cytosol. However, compared to Tau accumulations in sensor cells, recombinant full-length CFP-Tau showed less overall lifetime quenching even in heparin-induced fibrillar CFP-Tau aggregates, which might be accounted to differences between the cellular and the *in vitro* environment (level of crowding, ion composition, pH, presence of other biomolecules), but also to the fact that we compared TauRD (∼120 aa) in cells with full-length Tau (441 aa) *in vitro.* Compared to CFP-Tau, CFP-TauRD molecules are smaller and therefore have *per se* a higher density in condensates and a tighter packing of the CFP-Tag (=more quenching) in aggregates. Of note, it was shown that the fluorescent protein tag prohibits the packing of TauRD into dense amyloid-like aggregates (Kaniyappan *et al*, 2020), which may, however not be the case in Tau condensates.

One of the early changes observed during neuronal stress, in AD, and in tauopathies (Zempel & Mandelkow, 2014) is the missorting of Tau into the soma, which enables interactions of Tau with molecules and organelles not occurring in healthy neurons. Among these interactions is the accumulation of Tau at the cytosolic side of the nuclear envelope, which we observed in AD brain, as well as in a HEK cell model of templated Tau misfolding. Tau accumulations at the nuclear envelope occurred more often in cells treated with Tau condensates than with pre-formed Tau agrgegates (inducedwith heparin), opening the possibility that aged condensates harbor specific molecular Tau conformations or arragements that seed Tau condensation. Since nuclear envelope Tau can impair protein and RNA transport in and out of the nucleus (Eftekharzadeh *et al*, 2018; Cornelison *et al*, 2019), our findings provide a novel and intriguing link between Tau condensation and neurotoxicity in the brain.

## Abbreviations and Definitions

AD: Alzheimer’s disease
CFP: cyan fluorescent protein
DLS: dynamic light scattering
FTD: frontotemporal dementia
LLPS: liquid-liquid phase separation
MTs: microtubules
R_0_: radius of hydration (Stokes radius)
YFP: yellow fluorescent protein
FRAP: fluorescence recovery after photobleaching
FLIM: fluorescence lifetime imaging microscopy
FRET-FLIM: Foerster Resonance energy transfer measured by FLIM of the donor
CYT: cytosolic Tau inclusions
NUC: intranuclear Tau accumulations
NE: Tau accumulations at the nuclear envelope
hep_AGG_: Tau aggregated in the presence of heparin at a 4:1 (Tau:heparin) ratio
mesoscopic clusters: R_0_ ∼100-200 nm
microscopic condensates: R_0_ ∼1000 nm
monomeric/clusters/condensates: refers to degree of Tau assembling
Tau coacervation: LLPS driven by interaction of Tau with polyanions in the presence of PEG
Tau condensation: term for all LLPS processes

## Author contributions

J.H. performed most experiments, analyzed data, and helped preparing Figures. C.E. performed DLS experiments. M.F., A. D-B., L.D., S.F., H.B., and H.R. helped with experiments. M.K. and T.M. helped with FLIM imaging. S.K. provided antibodies and helped editing the manuscript. S.W. designed the study, S.W. and C.B.supervised experiments, S.W., C.B. and E.M. wrote and edited the manuscript and figures and provided the funding.

## Acknowledgements

We thank Eva Maria Mandelkow for data discussions, David Schwefel, Ulrike Kugelkorn, and Britta Eickholt at the Charité Berlin for their support in protein expression, and Sarah L Devos and Bradley Hyman for providing the AAV TauRD-CFP.P2a.TauRD-YFP construct. Microscopy, except FLIM, was performed at the AMBIO imaging facility of the Charité Berlin.

## Methods

### Human samples

Human AD (Braak 5/6) and age-matched control brain samples were received from the Biobanks at the Charité Berlin, Germany.

### AD brain lysate

AD brain lysates were prepared by homogenizing frozen frontal cortex tissue from AD brains with a dounce homogenizer in 3 volumes Tris-buffered saline (TBS) containing protease inhibitors (Pierce Protease Inhibitor Mini Tablets, EDTA-free). The homogenate was cleared by centrifugation at 10000 g for 10 min at 4°C, and the supernatant was aliquoted, frozen and kept at -80°C.

### Tau detection in AD brain sections

Paraffin brain sections of hippocampal CA1 were immunolabeled for phospho-Tau (PHF1, epitope pS396/pS404; AT8, epitope pS199/pS202/pT205; AT180, epitope pT231) followed by DAB deposition or immunofluorescence labeling as previously described (Eftekharzadeh *et al*, 2018). Sections were imaged in bright field using 40x or 63x objectives on a widefield fluorescence microscope (Eclipse-Ti, Nikon).

### Cloning of ECFP-Tau

To generate ECFP-Tau for bacterial expression, human full-length wildtype Tau (2N4R, hTau40) cDNA tagged with 6xHis at the C-terminus was introduced into a pNG2 vector (a derivative of pET-3a, Merck-Novagen) linearized with NdeI and BamHI, the ECFP cDNA, respectively, were introduced into the plasmid at the N-terminus using Gibson assembly, yielding the pNG2 ECFP-Tau-6xHis expression vector.

### Recombinant proteins

All plasmids were verified by Sanger sequencing prior to protein production. Human full-length wildtype Tau (2N4R, hTau40) and Tau^ΔK280^ were expressed in *E. coli* BL21 Star (DE3) (Invitrogen) and as previously described (Barghorn *et al*, 2005). Protein expression was induced with 0.5 mM IPTG at OD600=0.6 for ∼3 h at 37°C. For wild-type Tau, cells were harvested, resuspended in lysis buffer (20 mM MES, 1 mM EGTA, 0.2 mM MgCl2, 1 mM PMSF, 5 mM DTT, protease Inhibitors (Pierce Protease Inhibitor Mini Tablets, EDTA-free) and lysed using a French press. After initial purification by adding 500 mM NaCl and boiling at 95 °C for 20 min, cell debris was removed by centrifugation and the supernatant was dialyzed against low salt buffer (Buffer A: 20 mM MES, 50 mM NaCl, 1 mM MgCl2, 1 mM EGTA, 2 mM DTT, 0.1 mM PMSF, pH 6.8), filtered (0.22 µm membrane filter), run through a cation exchange column (HiTrap SP HP,5 ml, GE Healthcare), and eluted with a high salt buffer (Buffer B: 20 mM MES, 1000 mM NaCl, 1 mM MgCl2, 1 mM EGTA, 2 mM DTT, 0.1 mM PMSF, pH 6.8). Fractions containing Tau were pooled, concentrated using spin column concentrators (Pierce Protein concentrators; 10-30 kDa MWCO, Thermo Fischer Scientific), and run through a size exclusion column (Superose 6 10/300, GE Healthcare). Fractions containing purified monomeric Tau were concentrated as before and buffer exchanged to PBS, 1 mM DTT, pH 7.4.

His-tagged ECFP-Tau was expressed in *E. coli BL21 (DE3) Rosetta cells* following eth same protocol. Cells were harvested, resuspended in lysis buffer (50 mM Tris, 200 mM NaCl, 20 mM Imidazole, 10 % Glycerol, 0.1 % Triton X, 1 mM PMSF, 5 mM DTT and protease Inhibitors (Pierce Protease Inhibitor Mini Tablets, EDTA-free)) and lysed using a French press. After centrifugation (1 h at 35000 g at 4 °C), the supernatant of the lysate was filtered through a 0.22 µm membrane filter and loaded onto a Ni-NTA column (HiTrap, 5 ml, GE Healthcare) using a low imidazole buffer (Buffer C: 50 mM Tris, 200 mM NaCl, 20 mM Imidazole, 10 % Glycerol, 1 mM PMSF, 0.3 mM TCEP), and eluted with a high imidazole buffer (Buffer C: with 500mM Imidazole). Fractions containing were pooled, concentrated using spin column concentrators (Amicon Ultra 15 concentrators; 50 kDa MWCO, Millipore), and run through a size exclusion column (Superdex200 16/600, GE Healthcare) using PBS, 0.3 mM TCEP, pH 7.4 as running and final storage buffer.

Phosphorylated Tau (p-Tau) was expressed in insect (SF9) cells by infection with a recombinant baculovirus containing a pTriEx-Tau plasmid that encodes human full-length Tau (2N4R) with a N-terminal Strep-Tag (Schwefel *et al*, 2014). After cell lysis, the cleared lysate was applied to a 10 ml Strep-Tactin column (Iba) and a gel filtration column (Superdex200 16 600, GE Healthcare). The purified protein was stored in PBS containing 1 mM TCEP at -80°C. Final Tau concentrations were measured by BCA assay (BCA kit, Pierce) and the proteins were stored at -80°C.

### Fluorescent labeling of protein and RNA

Tau proteins were fluorescently labeled using amine-reactive DyLight488-NHS ester (Thermo Scientific) following manufacturer instructions. The dye was dissolved in DMSO to a final concentration of 10 µg/µl, then added to the protein in PBS with 1 mM DTT in a 5-fold molar excess, and the labeling reaction was done for 4h at room temperature. Excess dye was removed by dialysis (Pur-A-Lyzer Mini Dialysis tubes, Sigma-Aldrich; MWCO 12-14 kDa) against PBS, pH 7.4, 1 mM DTT at 4°C. The labeling degree (amount of dye per molecule protein) was determined by measuring the final protein concentration and correlating it to the maximum absorbance of the attached dye. For calculations see manufacturer instructions. PolyA-RNA was covalently labeled with Cy3 using a LabelT-Cy3 kit (Mirus Bio MIR 3625) following the manufacturer instructions.

### In vitro phosphorylation of Tau and Phos-tag^TM^ Gel

Tau (2N4R, htau40) was *in vitro* phosphorylated by incubating 5-6 mg/ml of protein in phosphorylation buffer (25 mM HEPES, 100 mM NaCl, 5 mg MgCl, 2 mM EGTA, 1 mM DTT, Protease Inhibitors), recombinant kinases (2500 U/mg protein for PKA (NEB-P600S), 500:1 Tau:kinase for Gsk3ß (BPS-40007), and Cdk5/p25 (Sigma-C0745)) and 1 mM ATP over night at 30 °C and 250 rpm. After addition of NaCl to a final concentration of 500 mM and boiling for 10 min at 95 °C, denatured kinases were spun down at 100000 g for 30 min. The phosphorylated Tau in the supernatant was dialyzed against 25 mM HEPES, 10 mM NaCl, 1 mM DTT, pH 7.4 and stored at -80 °C.

### Phos-tag^TM^ Gels

For the Phos-tag^TM^ (Fujifilm, Wako Chemicals) Gel analysis, 1 μg of different phosphorylated Tau variants was mixed with 6X reducing SDS-loading buffer, boiled at 95 °C for 5 min, and separated in 8 % Acrylamide Gels containing 50 µM Phos-tag^TM^ and ZnCl^2^ at 0.03 A for 4 h. The Gels were stained with SYPRO Ruby (Invitrogen) and visualization with a Typhoon scanner (Trio, GE Healthcare) using the 532 nm laser and a 610 nm band-pass filter.

### Microscopy of in vitro Tau condensed phases

Prior to all Tau LLPS experiments, the Tau storage buffer (PBS, 1 mM DTT) was exchanged against 25 mM HEPES, 10 mM NaCl, 1 mM DTT by dialysis. Tau LLPS was induced by adding 4.8-10 µg/ml polyA (Sigma; average length=200-6000 nucleotides (Deleault *et al*, 2005)), 4.8 µg/ml tRNA (Roche; average length ∼80 nucleotides), 2.8 µg/ml heparin (Applichem; MW=8- 25 kDa, ∼40-130 sugar moieties), or 5-10% (w/vol) PEG8000 (Promega; radius of gyration (*R_g_*) ∼3-5 nm (Cole & Ralston, 1994; Soranno *et al*, 2014)) to 5 µM Tau containing 1-3% of fluorophore labeled Tau-DyLight488. For imaging of Tau droplets, 2-3 μl of the solutions were pipetted on amine treated-glass bottom dishes (TC-treated Miltenyi, CG 1.5), or on glass dishes (MaTek P35G-1.5) treated overnight at room temperature with 10 µg/ml L-glutamate (Fluka) diluted in sterile ddH_2_O. Excess L-glutamate was removed by rinsing the dishes 3-times with sterile ddH_2_O. To avoid evaporation of the LLPS sample during microscopy, the imaging dishes were equipped with a ddH_2_O prewetted tissue lining the inner edges and closed. Imaging was performed on a widefield fluorescence microscope (Eclipse-Ti, Nikon) or on a spinning disk confocal microscope (Eclipse-Ti CSU-X, Nikon) using 60x objectives.

### Tau:tubulin co-condensation assay

Tau (25 μM Tau in 25 mM HEPES pH 7.4, 1 mM DTT; 1% Tau-Dylight488), or 60min-old Tau:RNA or Tau:RNA:PEG coacervates were mixed with 5 μM porcine tubulin-Alexa594 (Cytoskeleton), and samples were pipetted onto the glass surface of an imaging dish (MaTek) 30 min later. The emerging coacervates were imaged on a confocal spinning disc microscope (Leica) with simultaneous excitation of Tau-Dy488 and tubulin-A594.

### Tau:tubulin co-condensation and microtubule bundle formation

Tau (25 μM Tau in 25 mM HEPES pH 7.4, 1mM DTT containing 1% Tau-Dylight488) or 60 min-old Tau:polyA or Tau:PEG:polyA condensates containing 10% Cy3-labeled polyA were imaged prior and after the addition of 5 μM tubulin (bovine cycled tubulin, PurSolutions-PUR-032005) containing 10% HiLyte647-labeled tubulin (porcine tubulin, Cytoskeleton-TL670M). For tubulin bundle formation, 5 µM tubulin and 1mM GTP were added to Tau or Tau condensates (25uM, after 60 min) in 1x BRB80, 1mM DTT, pH 6.8. The emerging condensates and MT bundles were imaged on glass bottom imaging dishes (u-Slide Angiogenesis Glass bottom dishes, Ibidi) using a confocal spinning disc microscope (Eclipse-Ti CSU-X, Nikon) with a 60x objective.

### Turbidity measurements

Tau condensate samples (25-30 μl) were loaded into 384-well μClear plates (Greiner) and the absorbance at 600 nm was measured in a plate reader (Synergy H1, Biotek). Measurements were taken every 5 min for the duration of 1 h and then every 15 min for 2 h. To prevent sedimentation of the dense phase, the plate was shaken for 5 sec before each measurement. Tau alone and buffer samples were included as controls.

### Dynamic light scattering

For DLS experiments, solutions of Tau, tRNA and PEG8000 were prepared at a two-fold concentration and centrifuged for 15 min at 16000g at room temperature prior to DLS experiments. To induce Tau LLPS, RNA or PEG8000 were added to the Tau containing DLS cuvette in a 1:1 ratio, and quickly mixed by inverting the cuvette. DLS measurements and data acquisition were carried out with a Spectroscatter-301 (Xtal Concepts) equipped with a laser operating at a wavelength of 660nm. Sample solutions were measured at a fixed 90° scattering angle in quartz glass cuvettes (path length: 1.5 mm, Hellma Analytics) at 20°C. The viscosity was adjusted to 2.951cp for samples containing 5% PEG8000.

The obtained autocorrelation functions (ACFs; Expanded View Figure S1A) were averaged over 20 seconds for each data point. Averaged ACFs were fitted applying the CONTIN regularization method (Provencher, 1982), and the hydrodynamic radii, R_0_ (Expanded View Figure S1B), were calculated via the Stokes-Einstein relation Equation

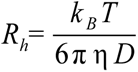

, with k_B_ being the Boltzmann constant, T the temperature, η the viscosity, and D the diffusion constant.

### Fluorescence recovery after photobleaching (FRAP)

Tau *in vitro* condensates containing ∼1% Tau-DyLight488 were imaged before and directly after bleaching (round ROIs; diameter ∼2-3 µm) with a 488 nm laser (90 % intensity; 3 loop), and the fluorescence recovery of the bleached region was measured for 40-60 s. For each bleached ROI a background ROI (outside condensates) and a reference-ROI (inside different condensates) of the same size were recorded in parallel. For FRAP of HEK293-TauRD^P301S^ cells, intracellular Tau accumulations were partially bleached (ROI diameter ∼1-2 µm) at a laser intensity of 70 % (2 loops) and the recovery of the bleached region was measured for 60 s. Background (outside cells) and reference signal (different cell) were recorded in parallel. For all FRAP measurements, the FRAP curves were background corrected and normalized to the background corrected reference signal. Experiments were performed on a spinning disk confocal microscope (Eclipse-Ti CSU-X, Nikon) using a 60x objective.

### Sedimentation assay

For sedimentation analysis of Tau condensates, 20 µl Tau LLPS samples were incubated for 1 h at room temperature and then centrifuged at 16000 g for 15 min to separate the light phase (in the supernatant) from the dense phase (in the pellet). After pipetting off 4 μl of the supernatant, the samples were centrifuged again at 16000 g for 5 min before 4μl of the pellet fraction were taken off, and same volumes of supernatant and pellet fractions were analyzed by SDS-PAGE.

### Tau LLPS and aggregation sample preparation

Condensate samples were prepared by adding final concentrations of 0.024 µg/ml tRNA (48:1; Tau:tRNA), 0.014 µg/ml heparin (82:1, Tau:heparin), or 5-10 % PEG8000 to 25 µM Tau (=1.15 mg/ml Tau) or Tau^ΔK280^ diluted in 25mM HEPES, pH 7.4, 1 mM DTT. Tau aggregation samples were prepared by adding final concentrations of 0.48 µg/ml tRNA (2.4:1, Tau:tRNA) or 0.287 µg/ml heparin (4:1, Tau:heparin) to 25 µM Tau. Samples were incubated at 37 °C for 1h, 24h, or 72h at 37°C. To prevent degradation of RNA, nuclease free H_2_O was used for all buffer preparations.

### SDS-PAGE, Semi-Native-PAGE, and Western blots

For SDS-PAGE, protein samples were mixed with 6X reducing SDS-containing loading buffer, boiled at 95°C for 5 min, and separated by SDS-PAGE (NuPAGE 4-12 % Bis-Tris, Invitrogen). For Semi-Native-PAGE, samples were mixed with a 6X loading buffer without SDS and reducing agents (BioLabs, B7025S) and without boiling loaded onto Native-PAGE gels (NativePAGE, Invitrogen). MES-buffer containing 0.1 % SDS (Invitrogen) was used as running buffer in both SDS-PAGE and Native-PAGE and 0.5 µg Tau were loaded per well. After separation, protein bands were stained with SYPRO Ruby (Invitrogen) and visualization with a Typhoon scanner (Trio, GE Healthcare) using the 532 nm laser and a 610 nm band-pass filter.

For Western blot analysis of phospho-Tau variants, 1.5-2ug total protein loaded were separated by SDS-Page, blotted onto a nitrocellulose membrane, which was then blocked with 3% non-fat dry milk in PBS at room temperature for 1 h and incubated with primary antibodies (rabbit anti-pS404 Tau, CST-20194S; rabbit anti-pS262 Tau, Invitrogen-44-750G; rabbit anti-pT205 Tau, Invitrogen-44-738G) diluted 1:1000 in blocking buffer (3% non-fat dry milk in PBS containing 0.05% of Tween (PBS-T)) overnight at 4°C. After washing with PBS-T and incubation with secondary Horseradish Peroxidase (HRP)-conjugated antibodies diluted 1:1000 in blocking buffer at room temperature for 1-2 h, the blots were developed using an ECL Western Blotting substrate (Promega) and a Fusion SL system (Vilber Lourmat). Afterwards, the blots were further incubated with a primary anti-total Tau antibody (rabbit Dako-A0024, 1:5000 in blocking buffer), then with the anti-rabbit HRP-conjugated secondary antibody, and developed again.

### Dot Blots

Tau LLPS and aggregation samples (5 µM Tau) were diluted to 0.5 µg protein in 50 µl using 25 mM HEPES, pH 7.4, 1 mM DTT and applied to a prewetted nitrocellulose membrane (GE Healthcare, 0.2 µm) under vacuum using a Dot Blot Apparatus (SCIE-PLAS). After loading, each well was washed twice with 100 µl PBS, and the entire membrane was blocked for 1 h with 3% BSA in PBS containing 0.05 % Tween (BSA/PBST). The membrane was incubated with primary antibodies (rat anti-oligomeric Tau 2B10, 1:250 (Chandupatla *et al*, 2020); rabbit anti-total human Tau antibody, Dako #A0024, 1:5000) overnight at 4°C, followed by incubation with secondary antibodies (anti-rat-800, LI-COR, 1:10000; anti-rabbit-680, Invitrogen, 1:10000) for 1-2 h at room temperature, and imaged with an Odyssey imager (LI-COR). Dot Blots were prepared in triplicates, and fluorescence intensity of each dot was measured using ImageJ and normalized to total Tau.

### Monitoring Tau seeding in vitro by Thioflavin-T fluorescence

Baseline aggregation of 10 µM Tau^ΔK280^ (0.46 mg/ml) was induced by adding 0.115 mg/ml heparin (4:1, Tau:heparin, [Tau^ΔK280^+hep]_AGG_). To determine the seeding potential of condensates, 0.0092 mg/ml Tau or Tau^ΔK280^ in form of 24 hour-old condensates (prepared with 25 µM Tau) were added to 10 µM Tau^ΔK280^ in PBS, pH 7.4, 1mM DTT (50:1, Tau^ΔK280^:condensates) and 50 µM Thioflavin-T (ThioT, Sigma-Aldrich) was added to detect Tau aggregation. [Tau^ΔK280^+hep]_AGG_ plus pre-aggregated Tau was used as a positive control. Tau^ΔK280^ in PBS without heparin and PBS with ThioT alone were used as negative controls. Samples were prepared in triplicates in clear bottom 384-well μClear plates (Greiner) and ThioT fluorescence (λ_Ex_=440 nm, λ_Em_=485nm) was recorded at 37°C in a plate reader (Synergy H1, Biotek) every 15 min after a 5 s shake.

### Cellular Tau seeding assay

HEK293 cells stably expressing the Tau-repeat domain (TauRD) containing the frontotemporal dementia (FTD)-mutation P301S and fused to CFP or YFP (HEK293 TauRD^P301S^-CFP/YFP (Holmes *et al*, 2014); ATCC #CRL-3275; cells were provided by Marc Diamond through Erich Wanker) were treated with OPTI-MEM culture medium containing 1.2 μM (0.055 mg Tau/ml) 24h- old Tau condensates for 2 h at 37°C. For seeding with AD brain lysate, cells were treated with 5 μg total protein. Prior to the treatment, Tau condensates were mixed with 0.8% lipofectamine 2000 (Invitrogen) to facilitate Tau uptake. After 2h, condensates and lipofectamine were diluted by addition of 3-volumens cell medium, and cells were incubated further for 24 h. Induced intracellular TauRD^P301S^ accumulations were detected by microscopy using the YFP fluorescence. Cells were imaged alive with a widefield fluorescence microscope (Eclipse Ti, Nikon) using a 20x objective for quantification of aggregates, or with a spinning disk confocal microscope (Eclipse-Ti CSU-X, Nikon) for morphological analysis and FRAP. Nuclei were counterstained with propidium iodide or DAPI. Quantification of the number of Tau accumulations per area covered by cells, and of different accumulation morphology types was done in Image J.

### Seeding of neurons expressing CFP/YFP-TauRD^P301S^

All procedures for experiments involving animals were approved by the animal welfare committee of Charité Medical University and the Berlin state government. Primary hippocampal mouse neurons were prepared as previously described (Hoffmann *et al*, 2019). Hippocampi of WT mice (C57BL/6J) P0–P2 brains were dissected in cold HBSS (Millipore), followed by 30 min incubation in enzyme solution (DMEM (Thermo Fisher Scientific), 3.3 mM cysteine, 2 mM CaCl_2_, 1 mM EDTA, and 20 U/ml papain (Worthington)) at 37°C. Papain reaction was inhibited by incubation of hippocampi in inhibitor solution (DMEM, 10% FBS (Gibco)), 38 mM BSA (Sigma-Aldrich), and 95 mM trypsin inhibitor (Sigma-Aldrich) for 5 min. Cells were triturated in Neurobasal-A medium (2% B27, 1% Glutamax, 0.2% P/S; (Thermo Fisher Scientific)) by gentle pipetting up and down. Isolated neuronal cells were plated at a density of ∼25,000 per 1 cm^2^ on μ-Slide 8-well culture dishes (ibidi) coated with poly-L-lysine and cultured in Neurobasal-A medium at 37°C and 5% CO_2_ until used for experiments. On day 3 *in vitro* (DIV3), the neurons were treated with AAV particles (AAV9 CFP-TauRD^P301S^.P2a.YFP-TauRD^P301S^) to induce the expression of equimolar amounts of CFP- and YFP-tagged TauRDP301S. On DIV6, the neurons were treated with AD brain lysate (TBS extract) to a final amount of 5 μg total protein per well. Six days later (at DIV12), the neurons were imaged in a brightfield microscope with a 60x objective.

### Probability of accumulations in cells

The probability of accumulation types in cells was determined from three confocal spinning disc images taken in three wells (8-well dishes) per condition. Since the seeding potential is different between the conditions (i.e. low seeding for Tau:PEG, high seeding for Tau:RNA and Tau:PEG:RNA, intermediate seeding for Tau aggregates; see Figure 6D), also the number of cells with accumulations (n*_cell_*) differs in the taken images.

In all cells having Tau accumulations, we determined whether the respective accumulation type (CYT, NUC, or NE) is present, regardless of the presence of multiple accumulations of the same type or the co-presence of other accumulation types. Most cells had more than one type of accumulations.

### Immunofluorescence labeling of cells

Cells were rinsed with PBS and fixed with 4% PFA in PBS for 10 min at RT, followed by two washes. In tris-buffered saline (TBS) for 5 min each. Cells were permeabilized with 0.3% Triton X-100 in PBS for 20 min at RT, followed by blocking with 3 % normal goat serum (NGS) in PBS for 60 min at RT and incubation with primary antibodies (mouse anti-RAN, 1:500, BD-610340, BD Biosciences; rabbit anti-laminB1, 1:1000, ab16048, Abcam; mouse anti-NPC, 1:1000, ab24509, Abcam; rabbit anti-POM121, 1:1000, NBP2-19890, Novus Biologicals) in 3% NGS in PBS overnight at 4°C. After 3 washes with PBS, cells were incubated with fluorescently labeled (Alexa-labeled antibodies, Invitrogen) secondary antibody diluted 1:1000 in 3% NGS in PBS for 1-2 h at room temperature. Nuclei were stained with DAPI diluted 1:1000 in PBS for 10 min. Cells were washed once more with PBS and stored at 4°C in PBS with 0.02% NaN3 until imaging.

### Fluorescent lifetime microscopy (FLIM) of ECFP-Tau condensates and HEK sensor cells

Tau:PEG:polyA condensates formed with 10 µM ECFP-Tau in 25 mM HEPES, 1 mM DTT, pH 7.4 were imaged 1 h or 24 h after formation. Tau aggregates were produced by incubating 60 µM ECFP-Tau with heparin (1:4 molar ratio of heparin:Tau) in PBS, 1 mM DTT with 0.02 % NaN3 and incubated at 37 °C for 6 days. For FLIM, 2-3 µl of each sample were pipetted on amine-treated glass bottom dishes (TC-treated Miltenyi, CG 1.5).

HEK293 TauRD^P301S^-CFP/YFP and HEK293 TauRD^P301S^-CFP cells treated with Tau condensates were fixed with 4% PFA in PBS prior to confocal FLIM on a Stellaris 8 FALCON (Leica) equipped with a 100x objective. Fluorescent intensities of CFP (λ_Ex_=430 nm) and YFP (λ_Ex_=510 nm) were recorded to select the ROI for FLIM. CFP fluorescent lifetimes were recorded at λ_Ex_=430 nm with a laser intensity of 4.5% to give at least 50000 photon counts per detected lifetime after decomposition. CFP lifetimes of soluble Tau, Tau condensates, and Tau accumulations in cells were defined in each individual FLIM image by placing circular ROIs in phasor plots of lifetime distributions, which enabled a fit-free lifetime decomposition. This analysis was done with the LAS X software (Leica). Reduced CFP lifetimes (LT-2, LT-3) indicated CFP:YFP FRET and/or CFP lifetime reduction due to dense packing.

### Data and statistical analysis

Image analysis was done in ImageJ. All data plotting, analysis, and statistical evaluation was performed using GraphPad Prism 8. Comparison of two groups was done by Student’s t-test, multiple groups were compared by one-way ANOVA with Tukey or Holmes-Sidak test for multiple comparison, as indicated in the figure legends. ****p<0.0001, ***p<0.001, **p<0.01, * p<0.0.5.

### Data Availability

All data is included in the manuscript and no data deposited in external repositories.

## Expanded View Figure Legends

**Expanded View Figure EV1.**
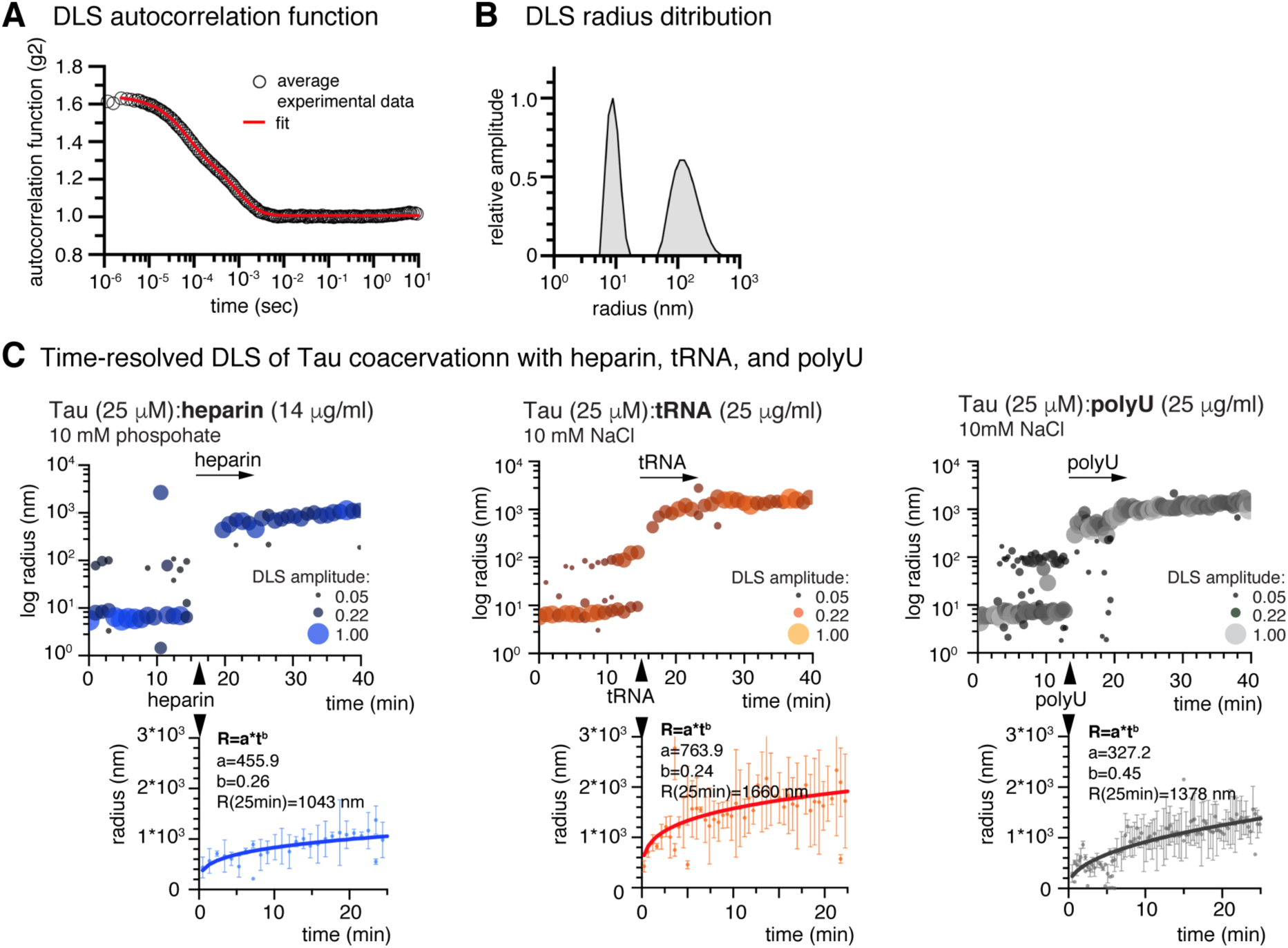
trDLS data of Tau coacervation. A Averaged experimental auto correlation function (black circles) and data fit applying CONTIN analysis (red line) of example trDLS data. Similar data was recorded and fitted for each 20 s interval of the trDLS measurements. B Distribution of hydrodynamic radii derived from autocorrelation data example in (A). C trDLS of Tau condensation upon addition of heparin, tRNA, and polyU RNA. Data point bubble sizes correspond to DLS amplitudes, which are proportional to the intensity of scattered light of the respective particles. Kinetics of condensate growth are assessed through power law, R=a*t^b^, fitting to trDLS data starting at the time of LLPS inducer addition. Data shown as mean±SD.

**Expanded View Figure EV2.**
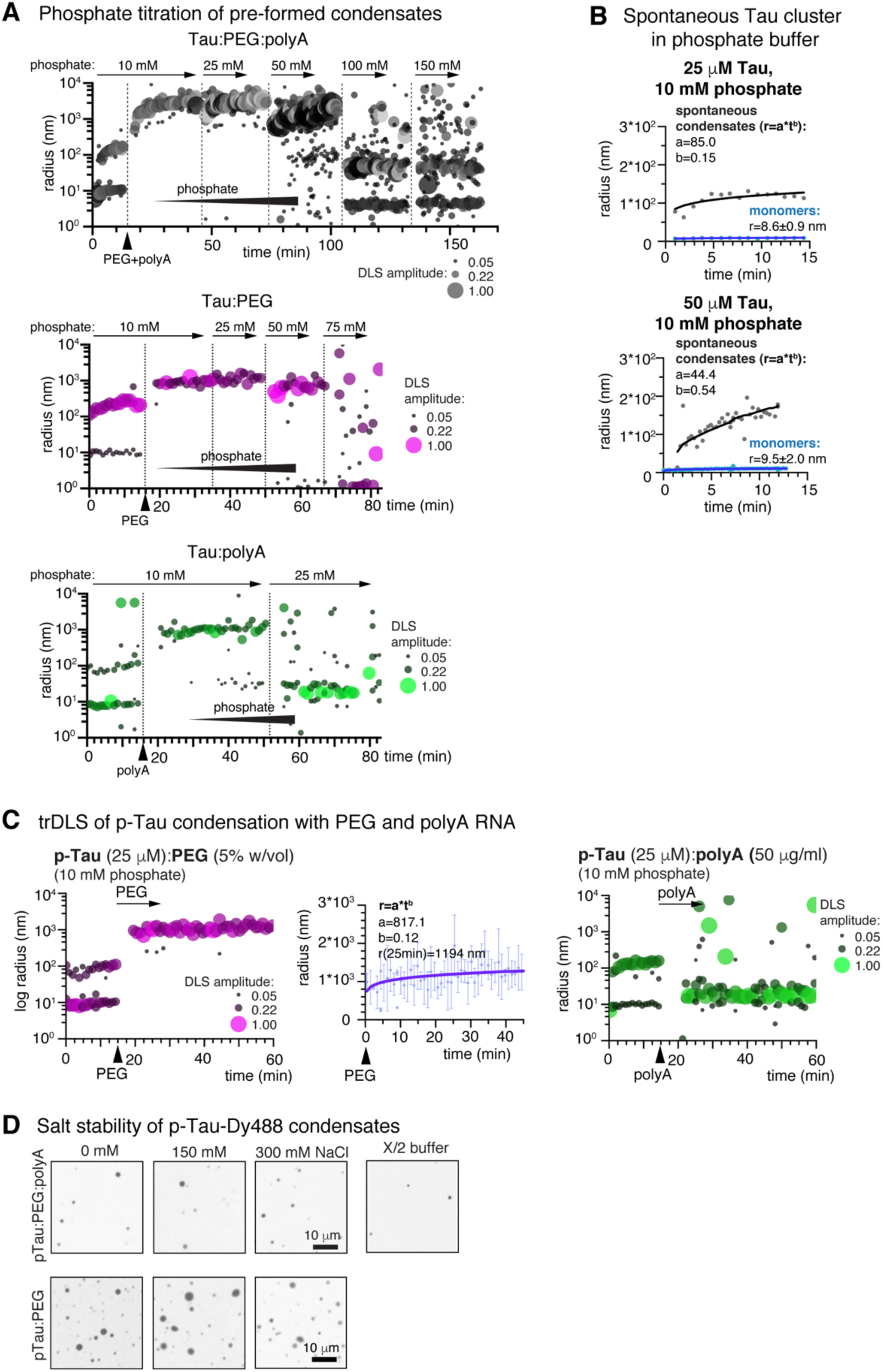
Effect of phosphate on Tau condensation. A trDLS of Tau:PEG:polyA, Tau:PEG, and Tau:polyA condensation, with a subsequent stepwise increase of phosphate ions. Tau:polyA condensates disappear at 25 mM phosphate ions in the buffer. Tau:PEG condensates dissolve at 75 mM, and Tau:PEG:polyA at 100 mM phosphate. Data point bubble sizes correspond to DLS amplitudes, which are proportional to the intensity of scattered light of the respective particles. Dashed vertical lines indicate times of phosphate addition. B Mesoscopic cluster growth at 10 mM phosphate in the buffer, at 25 μM and 50 μM Tau. Data were derived from trDLS measurements before the addition of LLPS inducers. Kinetics of cluster growth were assessed through power law, R=a*t^b^, fitting to trDLS data. C trDLS of p-Tau (from insect cells) condensation in p-Tau:PEG (left) and pTau:polyA (right) systems. Power law fit to condensate growth shows a low coarsening coefficient of b=0.12. polyA fails to induce coacervates of pTau. Data point bubble sizes correspond to DLS amplitudes, which are proportional to the intensity of scattered light of the respective particles. Kinetics of condensate growth were assessed through power law, R=a*t^b^, fitting to trDLS data starting at the time of LLPS inducer addition. Data shown as mean±SD. D Microscopy of pTau:PEG and pTau:PEG:polyA condensates at increasing NaCl concentrations and in X/2 buffer (only for pTau:PEG:polyA). Scale bars=10 μm.

**Expanded View Figure EV3.**
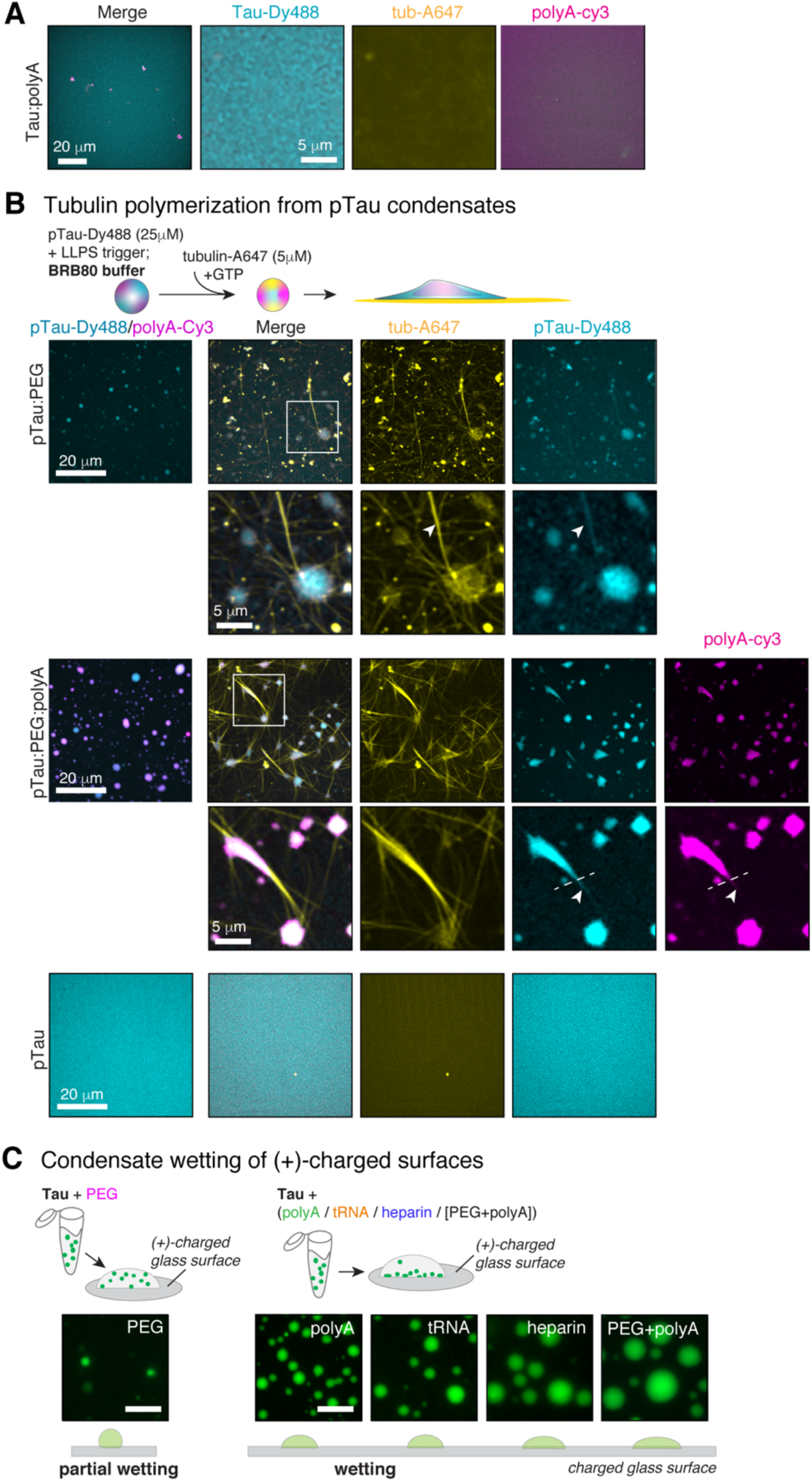
MT polymerization from p-Tau condensates. A Tau:polyA condensates do not form in MT polymerization buffer (BRB80). Scale bars=20 and 5 μm. B MT polymerization out of pTau:PEG and pTau:PEG:RNA condensates. Addition of HiLyte-647 (5 μM, 10:1 unlabeled:labeled tubulin) and GTP (1 mM) to p-Tau-Dy488 (25 μM) diluted in BRB80 polymerization buffer induces MT polymerization out of condensates. pTau-Dy488 alone did not induce MT polymerization. pTau:PEG systems show thin MT bundles, and pTau:PEG condensates attach and wet onto MTs. In pTau:PEG:RNA systems, we observe thick bundles attached to persisting pTau:RNA condensates, similar to Tau:PEG:RNA systems. p-Tau condensed phases extend along MT bundles beyond RNA condensed phase (indicated by dashed white line in inset), indicating preferred MT surface wetting by the Tau condensate. p-Tau alone does not promote tubulin polymerization. p-Tau was purified from insect cells. Images were taken 30 min after addition of tubulin. Scale bars=20 and 5 μm. C Attachment of Tau condensates to positively-charged glass surfaces (amine treated TC-dishes, Miltenyi). Tau:RNA and Tau:PEG:RNA condensates show surface wetting, whereas Tau:PEG condensates stay in solution or show partial wetting. Scale bars=10 μm.

**Expanded View Figure EV4.**
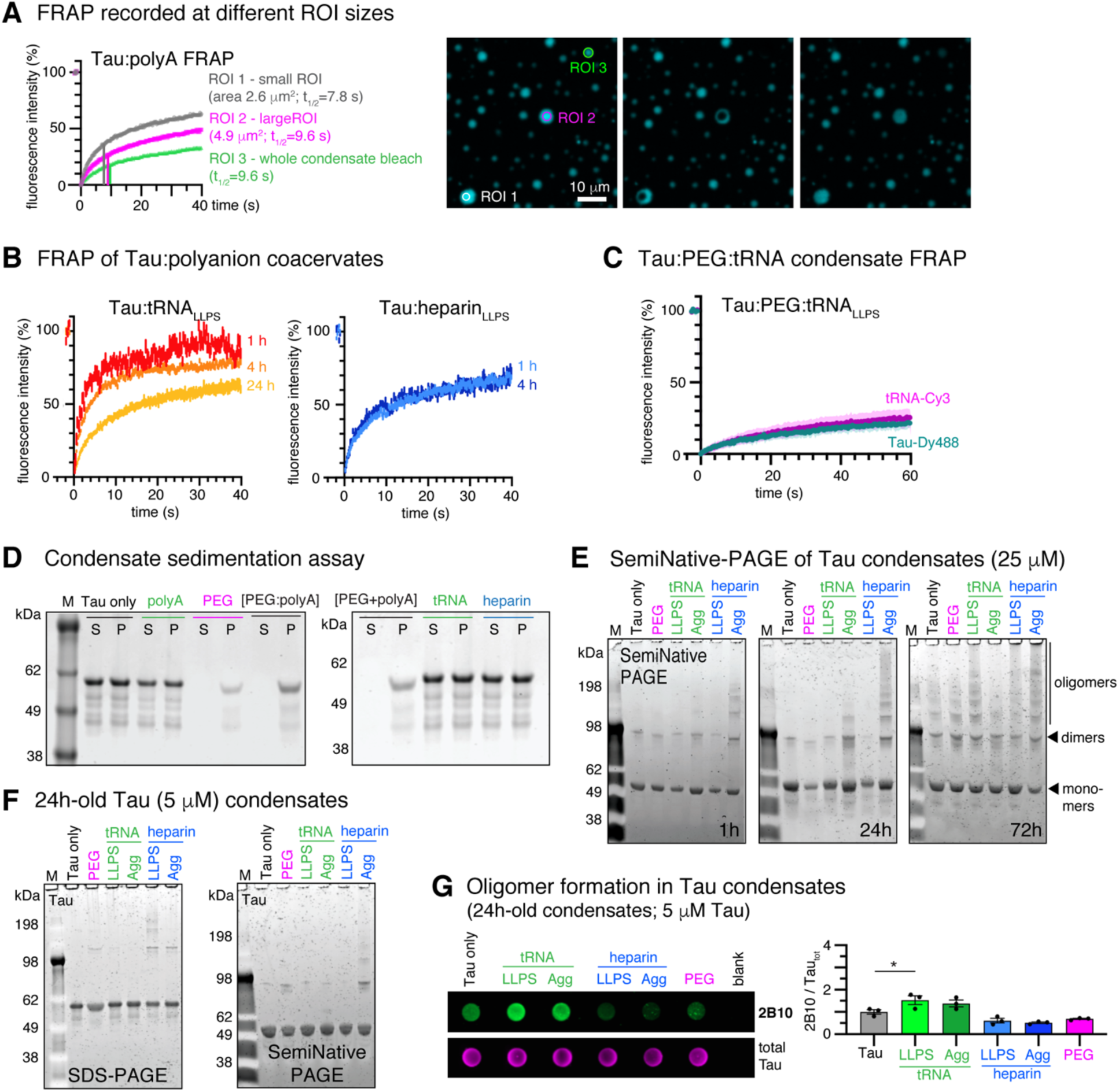
Percolation and oligomerization of Tau condensates. A FRAP of 4h-old Tau:polyA condensates comparing different FRAP ROI sizes at similar condensate size (ROI1 versus ROI2) and full condensate bleach (ROI3). Scale bar=10 μm. B FRAP of tRNA and heparin induced Tau coacervates show maintained molecular diffusion even after 4h and 24h. Data presented as mean±SEM, n=15 to 20 condensates per condition and time point. C Tau:PEG:tRNA condensates show little FRAP of Tau-Dy488 and tRNA-Cy3 after 4 h. Data shown as mean±SD. n=25 condensates per condition. D SDS-PAGE of Tau condensate sedimentation assay. Tau:RNA and Tau:heparin condensates do not enrich in the dense phase (P) upon centrifugation at 16000 g for 15 min at room temperature, hence have a lower density than Tau:PEG and Tau:PEG:polyA condensates, for which the dense (P) and light (S) phase can be separated by centrifugation. E Semi-Native PAGE of Tau incubated in LLPS (PEG, tRNA_LLPS_, heparin_LLPS_) or aggregation (tRNA_AGG_, heparin_AGG_) conditions for 1h, 24h, or 72h. Higher molecular weight Tau assemblies are prominent in aggregation conditions after 24h, and in all conditions after 72h, even in the Tau only control. Tau dimers are present already after 1h in all samples. F SDS-PAGE and Semi-Native PAGE of 5 μM Tau in condensation and aggregation conditions incubated for 24 h show the formation of SDS-stable Tau dimers in condensate. G Dot blot analysis of 24h-old Tau LLPS and aggregation samples indicates an increase of oligomeric Tau (2B10 antibody; p=0.043 for tRNA_LLPS_) in Tau LLPS samples containing tRNA or PEG compared to Tau only treatment. Data shown as mean±SD, n=3. One-way ANOVA with Holmes-Sidak test for multiple comparison.

**Expanded View Figure EV5.**
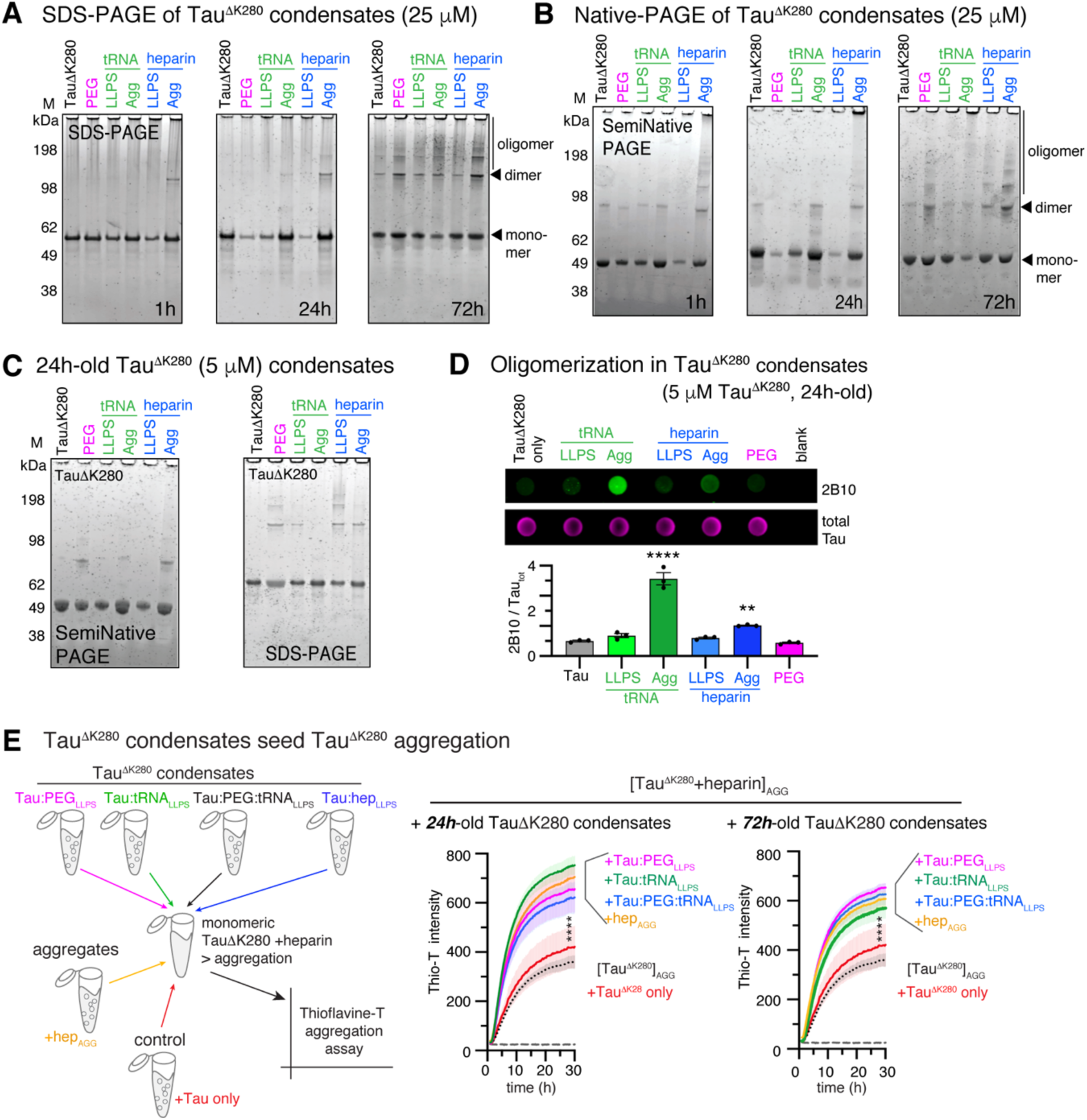
Oligomerization and seeding of pro-aggregant Tau^ΔK280^ condensates. A,B SDS-PAGE (A) and SemiNative PAGE (B) of Tau^ΔK280^ incubated in LLPS (PEG, tRNA_LLPS_, heparin_LLPS_) or aggregation (tRNA_AGG_, heparin_AGG_) conditions for 1h, 24h, or 72h. C Native PAGE and SDS-PAGE of 5 μM Tau^ΔK280^ in condensation and aggregation conditions incubated for 24 h. Even for low amounts of Tau^ΔK280^, PEG-, RNA- and heparin-induced condensates lead to the formation of SDS-stable Tau dimers. D Dot blot analysis of Tau^ΔK280^ oligomers (2B10 antibody) in 24h-old condensation and aggregation samples shows oligomeric Tau in aggregation conditions compared to Tau only treatment. Data shown as mean±SEM, one-way ANOVA with Holmes-Sidak post-test. E Thioflavin-T assay for the enhancement of *in vitro* Tau aggregation by Tau^ΔK280^ condensates. Tau^ΔK280^ (10 µM) incubated with heparin (molar ratio Tau:heparin=4:1; [Tau^ΔK280^]_AGG_) was used as base line condition for Tau aggregation. Addition of 24h-old or 72h-old Tau^ΔK280^ LLPS samples (9.2 μg/ml, molar ratio Tau^ΔK280^:condensates=50:1) to the reaction showed a significant increase of Tau^ΔK280^ aggregation for all LLPS conditions (PEG_LLPS_, tRNA_LLPS_, Petronel_las_), which was comparable to the effect of pre-aggregated Tau^ΔK280^ (hep_AGG_). Addition of Tau^ΔK280^ only showed no effect on Tau ^ΔK280^ aggregation. Data shown as mean±SD, n=3. One-way ANOVA with Tukey post-test for multiple comparison for values at 30 h.

**Expanded View Figure EV6.**
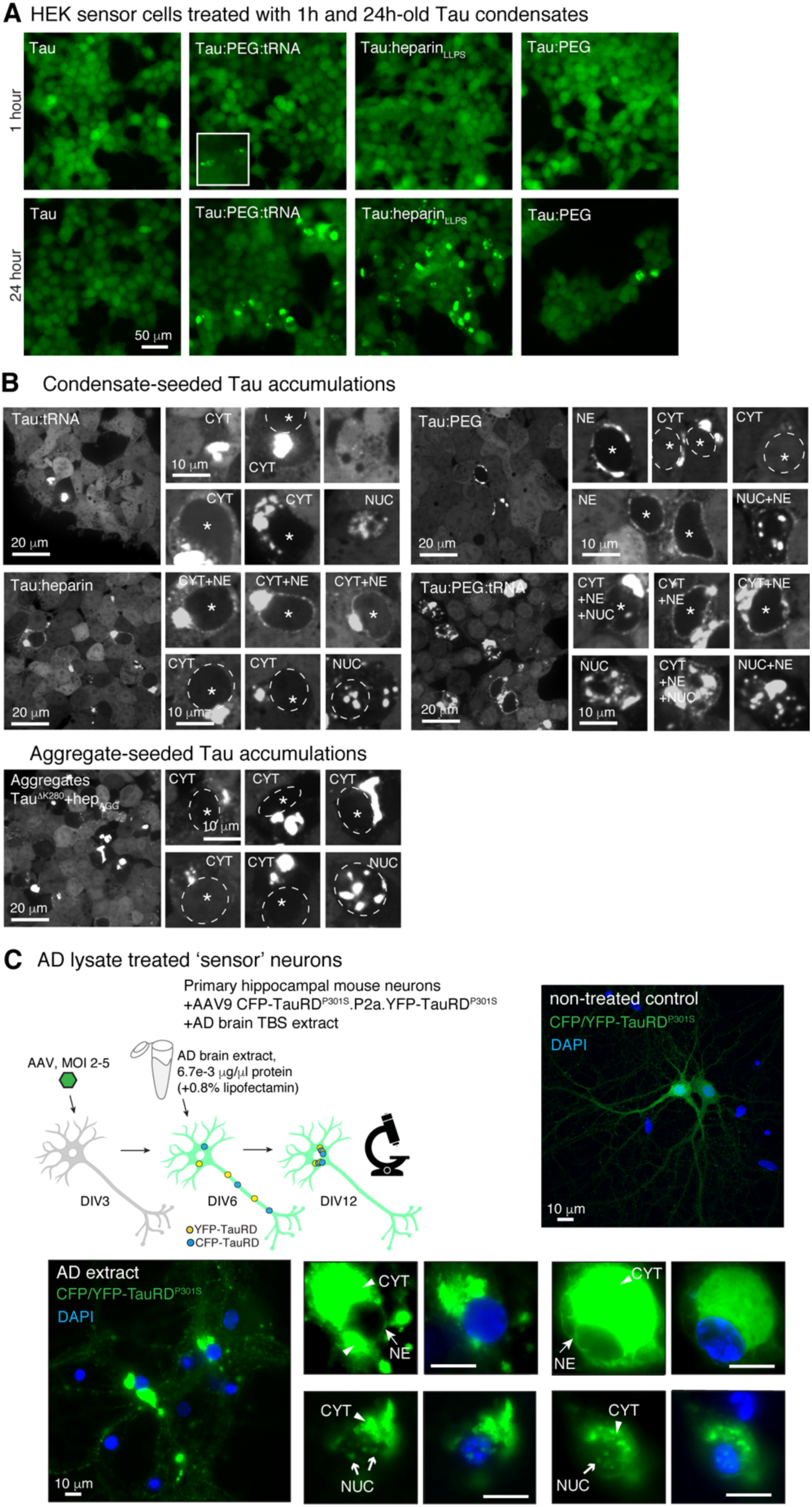
Tau accumulation “seeded” by 24h-old Tau condensates. A HEK sensor cells treated with 1h- and 24h-old Tau condensates. No seeding occurs in 1h-old LLPS preparations, whereas 24h-old condensates efficiently induce aggregation. Notably, a minor number of aggregates (inset) could be found already in 1h-old Tau:PEG:tRNA condensates, indicating a rapid production of seeding competent Tau in these condensates. B Confocal images of HEK sensor cells treated with Tau condensates (top) or Tau^ΔK280^ aggregates (bottom). Some nuclei are outlined with white dashed lines for better visualization. Scale bars=20 μm in overview, and 10 μm in close-ups. C Tau sensor neurons treated with AD brain lysate show Tau accumulations (CYT, NUC, and NE) similar to HEK sensor cells. Mouse primary hippocampal neurons were transduced with AAV CFP-TauRD^P301S^.P2a.YFP-TauRD^P301S^ and three days later treated with AD brain TBS extract. Tau accumulations were imaged 6 days later on DIV12. Non-treated neurons did not develop Tau accumulations. Scale bars=10 μm.

